# Targeting G_i/o_-coupled GPCRs to inhibit nociceptors: insights from the serotonin receptor Htr1b and triptans

**DOI:** 10.64898/2026.04.22.719367

**Authors:** Jing Peng, Brianna T. Sanchez, Anda M. Chirila, Xiangsunze Zeng, Michelle M. DeLisle, Lijun Qi, Jiayin Xiao, Karina Lezgiyeva, Sarah A. Low, Clifford J. Woolf, Nikhil Sharma, David D. Ginty

## Abstract

Pain perception is initiated upon activation of nociceptors of the dorsal root ganglia (DRG) and trigeminal ganglia. We identified G protein-coupled receptors (GPCRs) expressed in CGRP^+^ mouse and human DRG neurons and found that agonists of several identified G_i/o_-coupled and orphan GPCRs attenuated neuronal excitability. Experiments focusing on the G_i/o_-coupled serotonin receptor Htr1b, which is expressed in mouse and human CGRP^+^ DRG neurons, revealed that Htr1b/1d agonists, the triptans sumatriptan and zolmitriptan, attenuated CGRP^+^ neuron excitability *in vitro* and exhibited analgesia across several pain models, including neuropathic pain. Conditional genetic deletion experiments showed that triptan-induced analgesia is mediated by Htr1b expressed in A-fiber mechanonociceptors. Also, triptan-associated adverse effects are partially mediated by Htr1b-independent targets. Further testing identified the GPCR Gpr19 as an additional promising target for treating pain. These findings establish a preclinical screening platform for identifying novel analgesics and reveal nociceptor GPCRs that may be targeted to treat pain.

## Introduction

Detection of noxious stimuli and the perception of pain are essential for avoiding potential injury. However, pain, defined as “an unpleasant sensory and emotional experience associated with, or resembling that associated with, actual or potential tissue damage” by the International Association for the Study of Pain ^1^, can be debilitating and reduce the quality of life for many individuals. Current treatments for pain often lack efficacy and have adverse effects, and some, notably opioids, although effective, have contributed to the addiction epidemic due to their abuse liability. Because of the limited treatment options for pain disorders, there is an urgent clinical and societal need for understanding the underlying mechanisms of pain and developing new treatments for pain management.

The perception of pain is initiated by activation of a specific subset of primary sensory neurons called nociceptors whose cell bodies reside within the trigeminal or dorsal root ganglia (DRG) ^2^. These sensory neurons express many G protein-coupled receptors (GPCRs) ^3–5^. GPCRs are a large family of transmembrane proteins that respond to diverse extracellular stimuli and initiate a cascade of intracellular signaling events: the G_s_- and G_q/11_-coupled receptors upon activation generally promote neuronal excitation through stimulating adenylate cyclase (AC) and the formation of cyclic AMP (cAMP), or phospholipase Cβ (PLCβ), respectively. Conversely, G_i/o_-coupled GPCRs, such as the opioid receptors and metabotropic GABA receptors, can inhibit AC and stimulate G-protein activated Inwardly Rectifying K^+^ (GIRK) channels, thereby reducing neuronal excitability. A fourth subclass, the G_12/13_-coupled receptors are less well understood ^6^. Thus, activation of GPCRs in nociceptors can lead to either increased or decreased neuronal excitability, augmenting or diminishing the transduction of painful stimuli ^7–10^.

Many GPCRs serve as druggable targets for a range of therapeutics. Because of the unique role of G_i/o_-coupled GPCRs in reducing neuronal excitability, activating G_i/o_-coupled receptors may attenuate pain ^11,12^. Indeed, as the mainstream therapy against severe pain, morphine and other opioids target G_i/o_-coupled opioid receptors, preferentially the μOR ^13^. In addition, agonists of the G_i/o_-coupled serotonin receptors Htr1b/1d, the triptans, are a first line abortive treatment for migraine ^14^, and the G_i/o_-coupled α2-adrenergic receptor agonists, including clonidine and dexmedetomidine, are used clinically as an adjuvant in local anesthesia ^15,16^. Despite their broad clinical use, the cellular site of action of these medications is unclear.

Over the last decade, major progress has been made towards understanding the physiological and functional properties of somatosensory neuron subtypes, and recent transcriptomic approaches have revealed the genes expressed in these physiologically defined neuronal subtypes ^4,5,17–19^. Knowledge gained from these transcriptomic analyses has enabled us to generate genetic tools to manipulate select sensory neuron subtypes, including nociceptors ^5,18^. Moreover, DRG transcriptomic datasets can be used to identify GPCRs that are selectively expressed in nociceptors but not in other types of sensory neurons (i.e. low-threshold touch neurons and proprioceptors). These findings may be exploited to test the hypothesis that activation of inhibitory G_i/o_-coupled GPCRs that are selectively expressed in nociceptors reduce the sensation of pain following exposure to noxious stimuli, without affecting touch or proprioception.

Here, we utilized insights from somatosensory neuron transcriptomics combined with *in vitro* calcium imaging to identify candidate G_i/o_-coupled GPCRs expressed in mouse nociceptors that can be tested for their potential use for pain management. Several G_i/o_-coupled GPCRs were identified that when activated attenuate DRG nociceptor neuron excitability *in vitro*. Using Htr1b and the triptans as a proof-of-principle, we employed mouse genetics, electrophysiology, and a range of animal behavioral paradigms and found that activating this G_i/o_-coupled GPCR reduces the activity of nociceptors, attenuates transmitter release from nociceptor terminals, and reduces behavioral responses to noxious stimuli under normal conditions and in a neuropathic pain state. In addition, our findings reveal that an orphan GPCR, Gpr19, which we show to be G_i/o_-coupled, may also serve as a novel target for treating pain. We propose revisiting the therapeutic utility of existing and novel Htr1b agonists, beyond their use for treating migraine, and future identification and testing of agonists for Gpr19 and other GPCRs identified herein for their ability to treat different types of pain.

## Results

### Identifying GPCRs selectively expressed in mouse CGRP^+^ DRG sensory neurons

We identified GPCRs expressed in DRG somatosensory neuron subtypes implicated in pain signaling by assessing GPCR expression patterns using datasets of single-cell transcriptomic analyses of mouse DRG neurons ^5,18^. Prior functional analyses revealed physiological and morphological properties of transcriptionally defined somatosensory neurons, including A- and C-fiber low threshold mechanoreceptors (LTMRs), proprioceptors, C-fiber non-peptidergic heat-sensitive high threshold mechanoreceptors (*MrgprD^+^* and *MrgprB4^+^*C-HTMR/Heat neurons), C-HTMR/Heat/pruriceptors (*MrgprA3^+^* and *Sst^+^* C-HTMRs), cold-thermoreceptors (*TrpM8^+^*), and six transcriptionally distinct CGRP^+^ DRG neuron subtypes (Figure 1A). At least some CGRP^+^ DRG neuron subtypes may be considered nociceptors because they respond weakly or not at all to innocuous stimuli and strongly to noxious or painful stimuli ^18,20^. Our functional analyses have shown that these transcriptionally defined CGRP^+^ subtypes include two fast conducting A-fiber HTMR subtypes (*Bmpr1b^+^* and *Smr2^+^*Aδ-HTMRs), C-fiber noxious heat-sensing thermoreceptors (*Sstr2^+^* C-Heat), C-HTMR/Heat polymodal neurons (*MrgprA3^+^* C-HTMR/Heat), and two subpopulations with unknown physiological functions, CGRP-ε (*Oprk1^+^*) and CGRP-γ (*Adra2a^+^*) (Figure 1B) ^18,20^.

**Figure 1.**
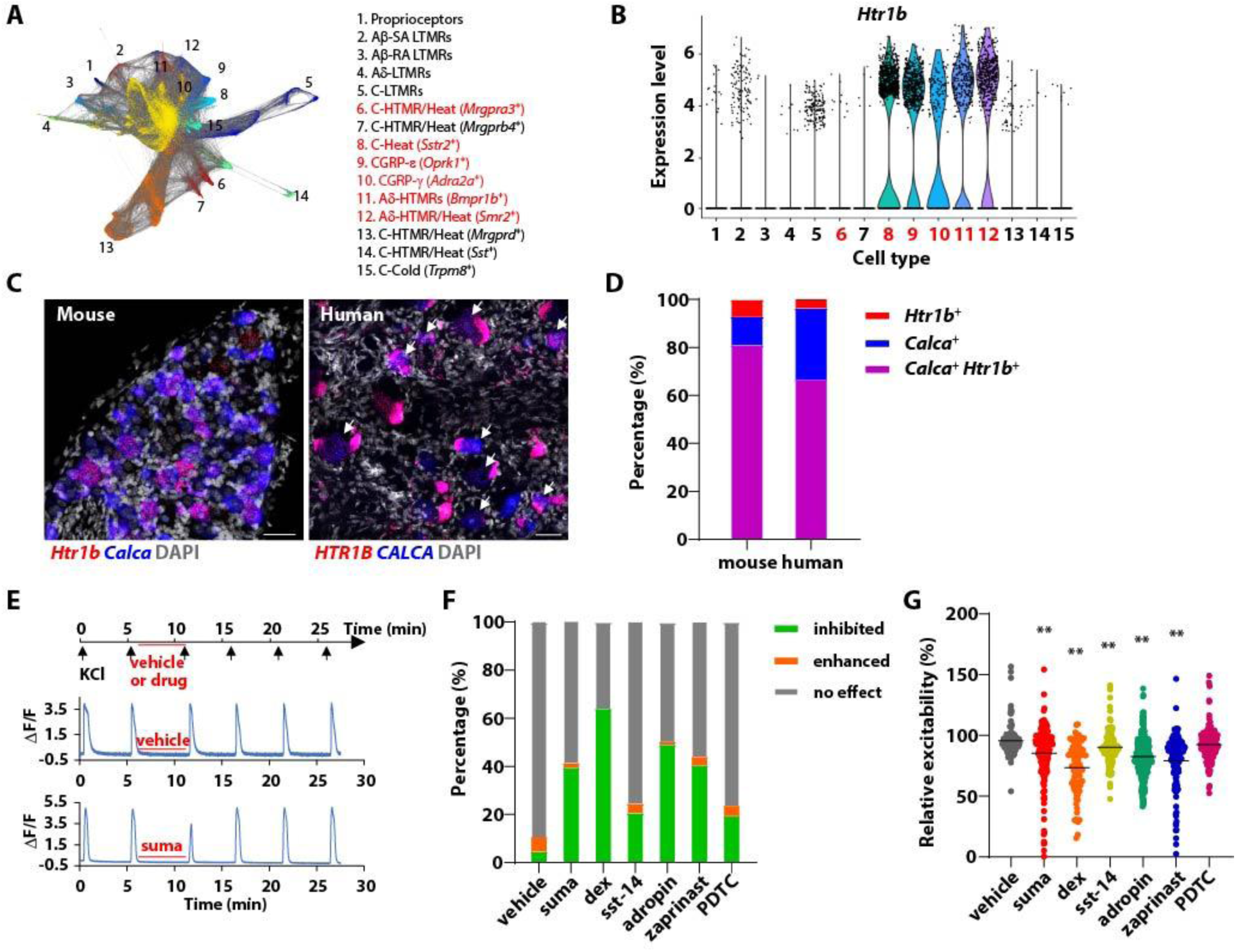
Activating G_i/o_-coupled GPCRs selectively expressed in CGRP^+^ DRG neurons, including the serotonin receptor Htr1b, decreases neuronal excitability. (**A**) Lay-out of sensory neuron subtypes in the mouse DRG from our published single cell RNA-sequencing dataset across development and into adulthood^5^. The physiological/functional property of each subtype is listed on the right. CGRP^+^ subtypes are highlighted in red. (**B**) The expression level of the serotonin receptor *Htr1b* in the mouse DRG sensory neuron subtypes. The cell types are number coded as in (A), with the CGRP^+^ subtypes highlighted in red. (**C**) *In situ* hybridization (RNAscope) confirmed co-expression of *Calca* and *Htr1b* in the mouse (left) and human DRGs (right). Human DRG cells co-expressing *CALCA* and *HTR1B* are indicated by arrows. Scale bar: 50 μm. (**D**) Quantification of (C). Percentages of cells that are positive or *Htr1b* only (red), *Calca* only (blue) or *Htr1b* and *Calca* (magenta) among all cells that are positive for at least one marker were plotted. (**E**) Top, diagram depicting the timeline of calcium imaging. Representative traces of cells treated with vehicle (middle) or sumatriptan (bottom). (**F**) A summary plot showing the percentage of cells inhibited or excited by the compounds tested, including the vehicle control (N=4), the Htr1b/1d agonist sumatriptan (10 μM, N=3), the α2-adrenergic receptor agonist dexmedetomidine (dex, 10 μM, N=4), the somatostatin receptor agonist Sst-14 (1 μM, N=2), and adropin (100 nM, N=4), zaprinast (30 μM, N=3) and PDTC (10 μM, N=3), the putative agonist of Gpr19, Gpr35 and Gpr149 respectively. (**G**) The tested compounds showed various degrees of inhibition of neuronal excitability. Each dot represents a cell.

To determine the expression profiles of murine GPCRs across somatosensory neuron subtypes, a curated list of over 800 non-olfactory GPCR genes ^21^ was mapped onto our single cell RNA-sequencing (scRNA-seq) dataset ^5^. This analysis revealed 69 GPCRs with expression patterns restricted to the CGRP^+^ sensory neuron types and not expressed in LTMRs or proprioceptors (Table S1). While some of these GPCRs are exclusively expressed in one of the CGRP^+^ neuron subtypes, others are more broadly expressed in several or all CGRP^+^ subtypes. Insights into heterotrimeric G protein transduction pathways used by the identified GPCRs were gathered using the IUPHAR/BPS Guide to Pharmacology. Among the GPCRs expressed in CGRP^+^ DRG neurons, several are known to be G_i/o_-coupled, including the serotonin receptor Htr1b (Figure 1B) and the α2-adrenergic receptor Adra2a (Table S1). Others, including Gpr19 and Gpr149, are orphan GPCRs whose transduction mechanisms are unclear (Table S1).

### GPCR expression patterns in human DRGs

Our long-term goal is to translate findings in mice into therapeutic opportunities for treating pain in humans. Therefore, we sought to identify those GPCRs with conserved expression patterns in mouse and human CGRP^+^ DRG neurons. We obtained acutely extracted human DRGs from a postmortem donor without a history of neurological disease and performed *in situ* hybridization (RNAscope) for the identified GPCRs. In human DRGs, *CALCA*, the gene encoding CGRP, was expressed in ∼70% of all neurons labeled by *NEFH* (NF200), which, unlike the mouse, serves as a pan-neuronal marker for human DRGs ^22^ (Figure S1A, B). This broad pattern of CGRP expression in the human DRG is consistent with previous findings ^23–27^. CGRP has been generally accepted and used as a marker for nociceptors in the human DRG studies ^23–27^. It is noteworthy that without functional validation, it is difficult to conclude that CGRP is a nociceptor-specific marker in human DRGs or whether the differences of CGRP expression between mouse and human DRGs translate into function. Parvalbumin (*PVALB*), a marker for mouse proprioceptors, is largely non-overlapping with CGRP in the human DRG (Figure S1C, D). Another marker, Somatostatin (*SST*), which is exclusively expressed in mouse C-HTMR/Heat neurons implicated in pruriception, is co-expressed with CGRP in the human DRG (Figure S1E, F), consistent with recent human DRG single-soma RNA sequencing data ^27^.

We examined the expression of a subset of candidate GPCR genes in the human DRGs. We prioritized known G_i/o_-coupled GPCRs that show restrictive expression in CGRP^+^ neurons in our mouse DRG scRNA-seq dataset, as well as orphan GPCRs with broad expression in mouse CGRP^+^ neurons, reasoning that some of the orphan receptors may be G_i/o_-coupled. The serotonin receptor, *HTR1F*, which is selectively expressed in mouse *Sst^+^* C-HTMR/Heat/pruriceptors, did not overlap with *SST* but rather with CGRP in the human DRG (Figure S1G, L). The α2-adregergic receptor *ADRA2A*, which is highly expressed in one of the murine CGRP^+^ subtypes, was co-expressed with CGRP in the human DRG (Figure S1H, L). The serotonin receptor *Htr1b* is expressed in most murine CGRP^+^ neuron subtypes, including the *Bmpr1b^+^* and *Smr2^+^*A-HTMRs, *Sstr2^+^* C-Heat thermoreceptors, and the two subtypes with unknown physiological properties, CGRP-ε (*Oprk1^+^*) and CGRP- γ (*Adra2a^+^*) (Figure 1B; Table S1). *HTR1B* exhibited a highly overlapping expression pattern with CGRP in the human DRG (Figure 1C, D).

Additionally, the orphan GPCRs, *GPR19*, *GPR35* and *GPR149* were co-expressed with CGRP in the human DRG (Figure S1I-L). Expression patterns of other identified GPCRs were also examined in the human DRG (Table S1). Taken together, we found GPCRs with similar expression patterns, i.e. co-expression with CGRP, in mouse and human DRGs, along with others with notable differences (Table S1). Candidates exhibiting co-expression with CGRP in both mouse and human DRGs were considered of interest, and several were included in subsequent screening steps.

### DRG GPCRs exhibit distinct G protein coupling

A few orphan GPCRs showed conserved co-expression with CGRP in mouse and human DRGs, including *Gpr19*, *Gpr35,* and *Gpr149* with unknown G protein transduction mechanisms. Since we were searching for inhibitory GPCRs that may potentially inhibit nociceptors, we attempted to determine whether the human orthologs of these orphan GPCRs are G_i/o_-coupled using an *in vitro* signaling assay that detects changes in intracellular cAMP levels in a heterologous system. As predicted, dexmedetomidine, a selective α2-adrenergic receptor agonist ^28^, dose-dependently reduced cAMP accumulation in *ADRA2A*-expressing but not vector-expressing HEK293 cells (Figure S2A), consistent with the G_i/o_-coupling specificity of ADRA2A ^29^. We failed to observe a dose-dependent decrease of cAMP levels following application of adropin, a putative agonist of GPR19 ^30,31^ (Figure S2B), or kynurenic acid or zaprinast, putative agonists of GPR35 ^32–34^ (Figure S2C, D). Pyrrolidine dithiocarbamate (PDTC), a putative agonist of GPR149 ^35^, appeared to reduce cAMP at higher doses in *GPR149*-expressing cells, but showed the same effect in vector-expressing control cells (Figure S2E). Although these negative findings do not support G_i/o_-coupling specificity, it is unclear whether any of these compounds are *bona fide* agonists of the respective GPCRs or if they are potent enough to activate the receptors in this assay. When expressed at high levels, GPCRs often exhibit ligand-independent constitutive activity ^36^, a phenomenon that may be exploited to determine the signaling pathway mechanisms employed by orphan GPCRs without prior knowledge of their ligands ^37–40^. Indeed, compared to vector-expressing controls, cells over-expressing the G_s_-coupled β2 adrenergic receptor, *ADRB2*, exhibited significantly higher levels of cAMP, and application of its agonist, isoproterenol (iso), failed to induce further cAMP accumulation, demonstrating agonist-independent constitutive G_s_-coupling activity (Figure S2F). In contrast, over-expression of *ADRA2A* significantly decreased cAMP levels, consistent with G_i/o_-coupling specificity (Figure S2F). Similarly, *GPR19* over-expression inhibited cAMP accumulation, suggesting constitutive G_i/o_-coupling activity (Figure S2G), consistent with previous reports ^30,41^. Over-expression of *GPR35* did not reduce cAMP levels (Figure S2H), while *GPR149* over-expression resulted in a moderate increase of cAMP compared to vector-expressing control cells, suggesting potential G_s_-coupling activity (Figure S2I).

### Activation of G_i/o_-coupled GPCRs reduces CGRP^+^ DRG neuronal excitability *in vitro*

We have identified several G_i/o_-coupled GPCRs, along with the orphan receptor GPR19 that is likely to be G_i/o_-coupled, that are restrictively expressed in CGRP^+^ DRG neurons. To determine whether activating these GPCRs inhibits DRG neuron excitability, we utilized an *in vitro* calcium imaging assay that uses KCl depolarization-induced calcium signaling as a surrogate measure for evoked neuronal excitability in acutely dissociated DRG neurons ^42^. For this, the calcium reporter GCaMP6f was selectively expressed in CGRP^+^ neurons using *Calca^CreER^* mice and a Cre-dependent GCaMP6f reporter allele (Ai148). *Calca^CreER^*; *Ai148* mice were treated with tamoxifen to promote Cre-mediated recombination and thus expression of GCaMP6f in CGRP^+^ DRG neurons. *In vitro* calcium imaging was then performed using cultured DRG neurons harvested from these mice. Calcium signals were elicited by depolarizing the sensory neurons with 25 mM KCl applied as 15 s pulses with 5 min inter-stimulus intervals (Figure 1E). The first two KCl pulses were used to determine the baseline excitability of the neurons. After the second pulse of KCl, agonists or putative agonists of candidate GPCRs were applied for 5 min before a third KCl pulse, which was then followed by a washout period that included two more pulses of KCl (Figure 1E, top). Representative calcium traces are shown in Figure 1E. In vehicle-treated control cells, KCl pulses elicited calcium influx with highly consistent amplitudes and durations (Figure 1E, middle). Pretreatment with sumatriptan (10 μM), a selective agonist of the G_i/o_-coupled Htr1b/1d receptors, significantly attenuated the KCl-evoked calcium response (Figure 1E, bottom). Approximately 40% of CGRP^+^ DRG neurons exhibited an attenuated calcium response when pretreated with sumatriptan (Figure 1F). The relative excitability of neurons treated with sumatriptan (magnitude of response normalized to baseline) was lower compared to vehicle-treated control neurons (Figure 1G).

As with sumatriptan, agonists of other G_i/o_-coupled receptors, including dexmedetomidine, an α2 adrenergic receptor agonist, and Sst-14, an agonist for somatostatin receptors, attenuated calcium responses in subsets of CGRP^+^ DRG neurons (Figure 1F, G). Adropin and zaprinast, putative agonists for Gpr19 and Gpr35 respectively, also decreased KCl-evoked calcium signals, while the putative Gpr149 agonist, PDTC, was without effect in our assay (Figure 1F, G). Thus, several agonists of G_i/o_-coupled GPCRs or putative G_i/o_-coupled orphan receptors attenuate excitability of subsets of CGRP^+^ DRG neurons.

CGRP^+^ DRG neurons are a heterogeneous population including unmyelinated small-diameter C-fiber neurons and myelinated medium- to large-diameter A-fiber neurons (Figure 1B). The calcium imaging findings showed that sumatriptan attenuated excitability of neurons with cell body diameters ranging from small to large (Figure S3A, left), consistent with the broad expression of *Htr1b* in CGRP^+^ C- and A-fiber DRG neurons (Figure 1B). Notably, sumatriptan-attenuated neurons had larger cell body diameters than sumatriptan-insensitive neurons (Figure S3A, left). While ∼30% of small diameter (< 25 μm) neurons were inhibited by sumatriptan, over 40% of medium diameter (25-35 μm) and 65% of large diameter (> 35 μm) neurons were inhibited (Figure S3A, right). These findings suggest that although sumatriptan can act on both C- and A-fiber CGRP^+^ neurons, medium to large diameter A-fiber CGRP^+^ neurons may be more responsive to the drug. In contrast to sumatriptan, the α2 adrenergic receptor agonist dexmedetomidine mostly inhibited small diameter neurons (Figure S3B), consistent with our finding that the α2A adrenergic receptor is expressed in small diameter C-fiber CGRP^+^ neurons (Figure S1H, left). Similarly, Sst-14 preferentially inhibited small diameter neurons (Figure S3C), in line with the restricted expression of somatostatin receptors (*Sstr1* and *Sstr2*) to C-fiber CGRP^+^ neurons (Sstr3, 4 and 5 are not expressed in the mouse DRG). We did not observe a clear trend of cell diameters for the neurons responding to adropin or zaprinast, the putative agonists of Gpr19 and Gpr35, respectively (Figure S3D-F).

In summary, we have performed a screen for mouse nociceptor-specific G_i/o_-coupled GPCRs by bioinformatically surveying the expression patterns of GPCRs in the mouse DRG scRNA-seq dataset and then validating expression of a subset of candidates in human DRGs. This screen revealed dozens of GPCRs expressed in CGRP^+^ sensory neurons, at least some of which show conserved expression patterns between mouse and human. Moreover, agonists of several G_i/o_-coupled GPCRs and orphan receptors attenuate excitability of subsets of CGRP^+^ neurons *in vitro*.

### Attenuation of behavioral responses to noxious cold by sumatriptan

We next focused on one of the G_i/o_-coupled receptors identified, Htr1b, and two of its known agonists, to test the hypothesis that pharmacologic activation of G_i/o_-coupled receptors in CGRP^+^ sensory neurons can attenuate nociception. We chose Htr1b because it is a well characterized G_i/o_-coupled GPCR with known agonists, the triptans, that are used clinically to treat migraine, although the mechanism and site of action of these drugs remain unclear. Therefore, using Htr1b and triptans as a proof-of-principle, we sought to test the hypothesis that activating nociceptor-specific G_i/o_-coupled GPCRs leads to silencing of nociceptors and attenuation of nociception, and a focus on Htr1b may help reveal the site of action of triptans and whether they can be used for indications other than migraine.

The triptans are prescribed to treat migraine and cluster headaches, and their utility in other types of pain has not been extensively tested clinically^14^. However, *Htr1b* expression is not limited to the trigeminal system, and, as our findings show, it is broadly expressed in CGRP^+^ DRG sensory neurons across all axial levels. Moreover, our *in vitro* findings suggest that sumatriptan can reduce excitability of CGRP^+^ DRG sensory neurons, and thus sumatriptan treatment may attenuate somatic pain by blocking nociceptor signaling. Therefore, we systematically tested the potential analgesic effects of sumatriptan in mice using a battery of pain behavior assays. In a cold plate behavioral assay, sumatriptan reduced nocifensive behavioral responses evoked by noxious cold at a dose of 300 μg/kg, delivered intraperitoneally (i.p.), which is comparable to the dose clinically used in human patients ^43^ (Figure 2A, B). Sumatriptan pretreatment led to a reduction in the number of behavioral bouts to noxious cold exposure and an increase in the latency to the first reactive behavior (Figure 2A, B). In contrast, sumatriptan was ineffective in reducing noxious heat responsiveness in a hot plate assay (Figure 2C, D), acute mechanical pain in a pinprick assay (Figure 2E, S4A, S4B), or inflammatory mechanical and thermal pain induced by intraplantar injection of zymosan (Figure 2F, G).

**Figure 2.**
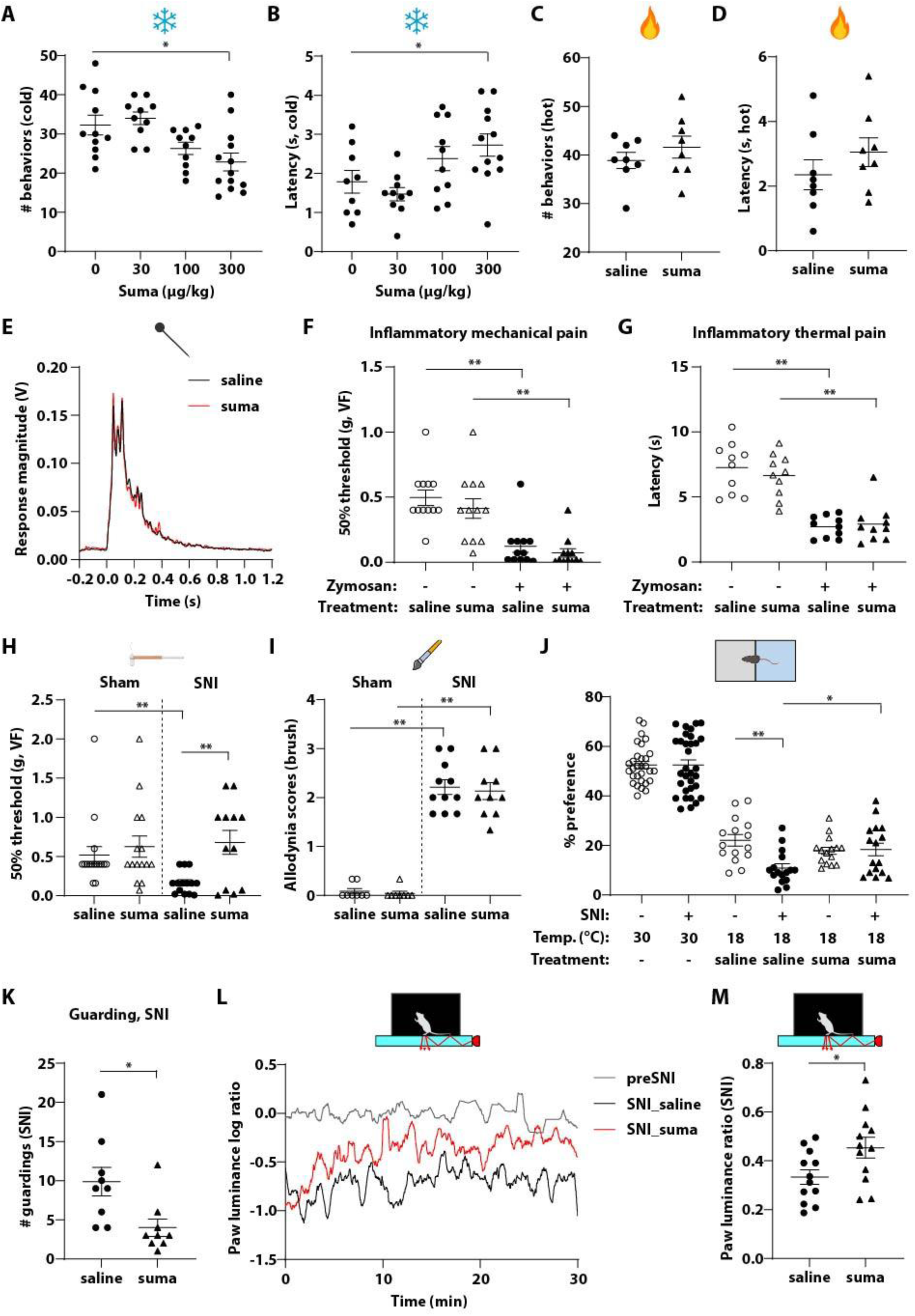
Sumatriptan attenuates responses to noxious cold and neuropathic pain in the spared-nerve injury (SNI) model. (**A, B**) Sumatriptan (30-300 μg/kg, i.p.) dose-dependently reduced nocifensive behaviors (A) and increased the latency to the first nocifensive behavior (B) in the cold plate (0°C) assay. The dose of 300 μg/kg was used for all other behavior tests unless otherwise specified. (**C, D**) Sumatriptan did not affect the number of behaviors (C) or latency (D) in the hot plate (51°C) assay. (**E**) Sumatriptan had no effect in reducing noxious mechanical pain in the pinprick assay. Averaged response traces of saline- (black) and sumatriptan-treated (red) animals were shown. Time 0 indicates the onset of the pinprick stimulus. (**F**) Inflammatory mechanical pain induced by intraplantar injection of zymosan (100 μg in 20 μL) was assessed by the von Frey (VF) test. The significant decrease of mechanical threshold after zymosan treatment was not alleviated by sumatriptan. (**G**) Inflammatory thermal pain was tested using the Hargreaves assay. Sumatriptan had no effect on zymosan-induced thermal hyperalgesia. (**H**) Sumatriptan alleviated punctate mechanical allodynia after SNI surgery in the von Frey test. (**I**) Sumatriptan did not affect the allodynia score in the dynamic brush test (C). (**J**) Sumatriptan reversed the cold allodynia after SNI surgery in the temperature preference assay. (**K**) Sumatriptan also attenuated guarding behaviors in SNI animals. (**L**) Representative traces of the paw luminance log ratio (calculated as logarithmic 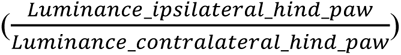 of preSNI (gray) and saline- (black) or sumatriptan-treated SNI animals (red) in the paw luminance assay. (**M**) Sumatriptan partially reversed the reduction of paw luminance ratio after the SNI surgery.

### Attenuation of neuropathic pain by sumatriptan

Next, we asked whether sumatriptan is effective in mitigating neuropathic pain. The effects of sumatriptan in non-cranial neuropathic pain have not been systematically investigated clinically, although there have been reports with mixed results in different preclinical models of neuropathic pain ^44–47^. We used the Spared Nerve Injury (SNI) model, which induces a robust and persistent form of neuropathic pain ^48,49^. Static/punctate allodynia and dynamic mechanical allodynia were tested in mice that underwent SNI surgery using von Frey (VF) filaments and dynamic brush, respectively. Sumatriptan reduced punctate mechanical allodynia, as indicated by reversal of the 50% response threshold in the VF assay (Figure 2H), however dynamic brush mechanical allodynia was unaffected (Figure 2I, S4C).

To assess cold hypersensitivity in the SNI model, we used a temperature preference assay in which animals were allowed to freely move in a chamber with the two sides of the floor set at 30°C/30°C or 30°C/18°C. Before SNI surgery, the mice showed no side bias when both sides of the floor were set at 30°C (∼50% preference). When given a choice between floor temperatures of 30°C or 18°C, control mice exhibited a preference for the 30°C side (Figure 2J). After SNI, animals showed no side bias at 30°C/30°C but did exhibit a more pronounced aversion to 18°C in the 30°C/18°C configuration (Figure 2J), suggesting that they had developed cold hypersensitivity. Interestingly, while sumatriptan did not alter temperature preference before the SNI surgery, sumatriptan given after SNI restored the cold preference of SNI mice to a level similar to that observed pre-SNI (Figure 2J). These findings suggest that sumatriptan attenuates cold hypersensitivity in a neuropathic pain model.

Two weeks after SNI, mice exhibited a spontaneous guarding behavior on the ipsilateral hind paw. As this behavior was observed during the animal’s resting state in the absence of external triggers, we speculate that it reflects spontaneous pain. This guarding behavior was attenuated by gabapentin, a medication commonly prescribed to treat neuropathic pain ^50^ (Figure S4D), suggesting this behavior reflects an aberrant sensory and not motor function. Similarly, sumatriptan significantly reduced the guarding behavior in SNI mice (Figure 2K).

A paw luminance assay that measures luminance-based paw surface contact using frustrated total internal reflection technology (FTIR) and simultaneously records the whole-body pose of the animals has also been used to detect spontaneous pain in a variety of pain models ^51^. Therefore, to further assess sumatriptan’s effect on spontaneous/ongoing pain, we tested its effect on SNI mice using the paw luminance assay. After SNI, animals exhibited altered weight bearing, i.e. less force applied to the ipsilateral (injured) hind paw, resulting in a reduced paw luminance ratio (ipsilateral to contralateral hind paw) (Figure 2L, S4E). Administration of sumatriptan attenuated the reduction of paw luminance ratio in SNI animals (Figure 2L, 2M,

S4E). An increase of paw luminance ratio was observed in both the locomoting and non-locomoting states (Figure S4F, G). The SNI surgery causes motor impairment and altered gait, which may confound the effects of sumatriptan in the change of paw luminance ratio. The simultaneous recording of whole-body pose in this assay allowed us to assess whether sumatriptan affected motor or gait of the SNI mice. There was no significant difference in travel distance, locomoting time, or mean speed between the saline- and sumatriptan-treated SNI mice (Figure S4H-J), ruling out a general effect on motor behavior. Moreover, the angle between the ipsilateral hind paw and the trunk and the distance between the hind paws were increased after SNI, but there was no difference between the saline- and sumatriptan-treated animals (Figure S4K, L), suggesting that sumatriptan did not alter the gait of the SNI animals. Therefore, it is unlikely that the increase of the luminance ratio observed in sumatriptan-treated mice was due to altered ambulatory behavior or gait.

Collectively, these findings suggest that the anti-migraine drug sumatriptan confers analgesia in somatic pain models. Sumatriptan reduced responses to noxious cold, although it was ineffective in other nociceptive pain assays, including noxious heat, mechanical and inflammatory pain (Figure 2A-G). Strikingly, sumatriptan exhibited an analgesic effect in the SNI model of neuropathic pain where it caused a reduction in static mechanical allodynia, cold hypersensitivity, and two measurements of spontaneous pain (Figure 2H-M).

### Sumatriptan acting through Htr1b reduces both CGRP^+^ neuron excitability and synaptic transmission in the spinal cord

We next sought to determine the potential sites of action of sumatriptan. Sumatriptan is a potent agonist for both Htr1b and Htr1d but exhibits a much lower affinity for the other serotonin receptors ^52–54^. Our sequencing data suggested that Htr1d is expressed at low levels in proprioceptors and C- and A-LTMRs (Figure S5A), while Htr1b is exclusively and highly expressed in CGRP^+^ DRG neurons (Figure 1B). We confirmed by RNAscope that *Htr1d* and *Htr1b* are expressed in a non-overlapping manner in both mouse and human DRG (Figure S5B, C). Since GCaMP6f was exclusively expressed in CGRP^+^ neurons in our *in vitro* calcium imaging assay, we speculated that the inhibitory effect of sumatriptan is mediated through Htr1b. To test this, we generated conditional knock-out (cKO) mice using *Calca^CreER^* and a Cre-dependent *Htr1b* conditional allele (*Htr1b^fl^*) ^55^ to selectively delete *Htr1b* in CGRP^+^ neurons that express GCaMP6f (*Calca^CreER^; Ai95; Htr1b^fl/fl^*, hereafter denoted *Htr1b* cKO*^Calca^*). Efficient deletion of *Htr1b* in CGRP^+^ DRG neurons was confirmed by RNAscope (Figure S5D, E). In *Htr1b* cKO*^Calca^* mice, sumatriptan failed to attenuate excitability of CGRP^+^ neurons (Figure 3A, B), indicating that Htr1b mediates the inhibitory effect of sumatriptan on CGRP^+^ DRG neuron excitability.

**Figure 3.**
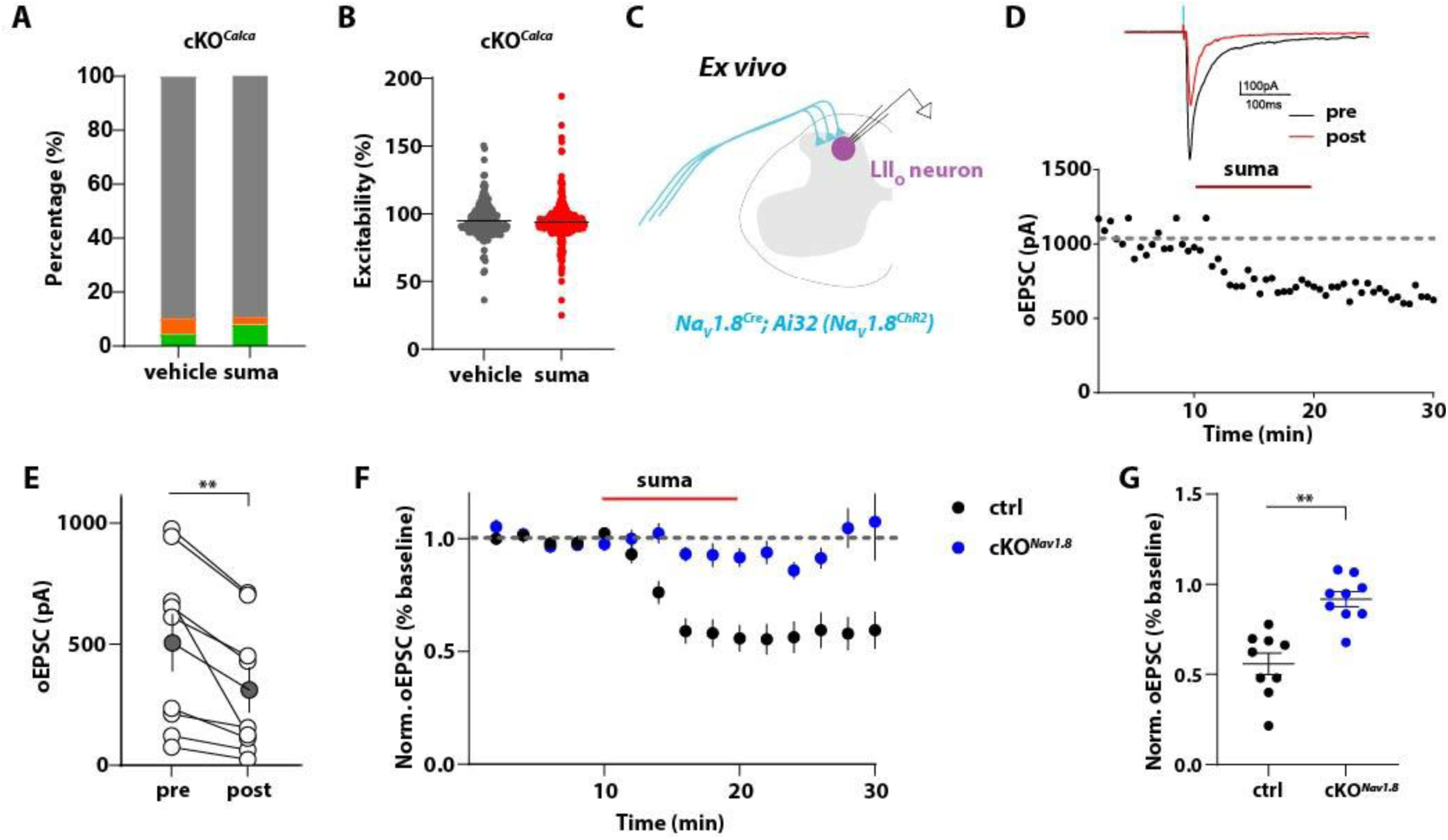
Htr1b in the sensory neurons is responsible for the inhibition of CGRP^+^ DRG neurons *in vitro* and the suppression of glutamate release from sensory neuron synapses in the spinal cord dorsal horn. (**A**) CGRP^+^ cells no longer respond to sumatriptan in *Htr1b* cKO*^Calca^* animals (N=3). (**B**) Vehicle- and sumatriptan-treated cells show comparable excitability in *Htr1b* cKO*^Calca^* animals. (**C**) Schematic diagram of the slice electrophysiology experiment. ChR2 was expressed in CGRP^+^ and other C-fiber sensory neuron subtypes using *Na_v_1.8^Cre^* and the Cre-dependent reporter Ai32. (**D**) Response of a representative lamina II dorsal horn neuron to sumatriptan bath-application. Sumatriptan application decreased the amplitude of optically-evoked EPSC (oEPSC). (**E**) Quantification of the oEPSCs before- and after- sumatriptan treatment (N=3, n=9). Each dot represents one neuron; gray dots show average oEPSC values. (**F**) Sumatriptan-mediated inhibition of oEPSC inhibition in control mice (black, N=3, n=9) was diminished in *Htr1b* cKO*^Nav1.8^*littermates (blue, N=4, n=9). (**G**) Quantification of (D).

The *in vitro* calcium imaging experiments indicate that sumatriptan attenuates neuronal excitability when applied directly to CGRP^+^ DRG neurons. *In vivo*, sumatriptan could act on cell bodies, peripheral axons, and/or central axon terminals of these sensory neurons. As a relatively hydrophilic compound, sumatriptan was initially considered unable to cross the blood brain barrier (BBB). However, recent findings have suggested that sumatriptan can cross the BBB and activate Htr1b/1d receptors in the central nervous system ^56–59^. To explore the possibility of a central site of action of sumatriptan in blocking nociceptor function, we performed *ex vivo* slice physiology in the spinal cord. We used *Na_v_1.8^Cre^*, which labels all CGRP^+^ neurons and additional C-fiber neuron subtypes ^60,61^, Cre-dependent channelrhodopsin (Ai32), and the *Htr1b^fl^* allele, to express the photoactivatable opsin ChR2 (*Na_v_1.8^ChR^*^2^) in *Htr1b^+/+^* and *Htr1b* cKO*^Nav1.8^* (*Na_v_1.8^Cre^; Ai32; Htr1b^fl/fl^*) mice. We prepared acute lumbar spinal cord slices from *Htr1b^+/+^* and *Htr1b* cKO*^Nav1.8^*mice and made whole cell patch clamp recordings from neurons within lamina I-II_o_ of the dorsal horn, where peptidergic C-fiber DRG neurons and some Aδ-HTMR fibers terminate ^18,20^ (Figure 3C). Excitatory post-synaptic currents (EPSCs) were evoked optogenetically by stimulating *Na_v_1.8^ChR2^*primary afferent axonal terminals with 1 ms pulses of blue light (473 nm, 5 mW) delivered at 30 s intervals. After a 10 min stable baseline, sumatriptan (10 μM) was then bath-applied for 10 min. We observed a significant reduction of optically evoked EPSC amplitudes (oEPSCs) following sumatriptan application in *Htr1b^+/+^* wildtype animals (Figure 3D, E). In *Htr1b* cKO*^Nav1.8^*mice (*Na_v_1.8^Cre^; Ai32; Htr1b^fl/fl^*), *Htr1b* was efficiently depleted, as verified by RNAscope (Figure S5F, G). Importantly, the attenuation of the oEPSC amplitude by sumatriptan observed in the *Htr1b^+/+^* control littermates was not observed in *Htr1b* cKO*^Nav1.8^* mice (Figure 3F, G), indicating that sumatriptan can act presynaptically through Htr1b receptors expressed in sensory neurons to reduce synaptic transmission. This finding is consistent with previous findings that sumatriptan can inhibit synaptic neurotransmission in trigeminal sensory neurons ^62–64^. These findings suggest that sumatriptan attenuates both excitability and neurotransmitter release in CGRP^+^ DRG neurons through its receptor Htr1b.

### Generating and characterizing mice lacking Htr1b in adult somatosensory neurons

The *in vitro* and *ex vivo* findings suggest that Htr1b expressed in sensory neurons is responsible for triptan-mediated inhibition of CGRP^+^ DRG neuron excitability and suppression of presynaptic neurotransmission in the spinal cord. To determine whether the analgesic effects of sumatriptan are mediated by Htr1b and/or Htr1d, or some other target, and to test whether somatosensory neurons are its primary site-of-action, we generated mice that lack Htr1b receptors in all somatosensory neurons using a pan-somatosensory neuron CreER driver line (*Avil^CreER^; Htr1b^fl/fl^*, herein denoted *Htr1b* cKO*^Avil^*). CreER-mediated recombination was induced by exposing young adult mice (P28-32) to tamoxifen (1 mg/day for five days) to excise *Htr1b* in somatosensory neurons. Knock-out efficiency in DRG neurons was ∼74%, as determined by RNAscope (Figure S6A, B).

The pan-sensory neuron *Htr1b* cKO*^Avil^* mice appeared grossly normal compared to control littermates, with comparable body size, weight and viability. They also exhibited normal locomotor activity in the open field (OF) test (Figure S6C-D). Mechanical and temperature sensitivity of *Htr1b* cKO*^Avil^* mice were also comparable to their control littermates (Figure S6E, F). In a sandpaper avoidance behavioral test, while control animals consistently avoided a rough texture, *Htr1b* cKO*^Avil^*mice lost this preference (Figure S6G). Taken together, these findings suggest that *Htr1b* cKO*^Avil^* mice exhibit grossly normal behaviors, except for an impaired avoidance of an aversive texture.

### Analgesic actions of sumatriptan are mediated through Htr1b expressed in somatosensory neurons

We used the *Htr1b* cKO*^Avil^* mice to test the possibility that sumatriptan acts on Htr1b expressed in somatosensory neurons to promote its analgesic effects. Indeed, in the noxious cold plate assay, sumatriptan reduced nocifensive behaviors in control animals (Figure 4A, B), as predicted from our prior findings (Figure 2A, B), however it failed to do so in *Htr1b* cKO*^Avil^*mice (Figure 4A, B). Both the magnitude and latency of responses to noxious cold were unaltered by sumatriptan in *Htr1b* cKO*^Avil^* mice. Similarly, in the SNI model of neuropathic pain, sumatriptan reduced the guarding behavior observed in control but not *Htr1b* cKO*^Avil^*animals (Figure 4C). These findings indicate that the analgesic effects of sumatriptan are mediated by Htr1b receptors expressed in the sensory neurons.

**Figure 4.**
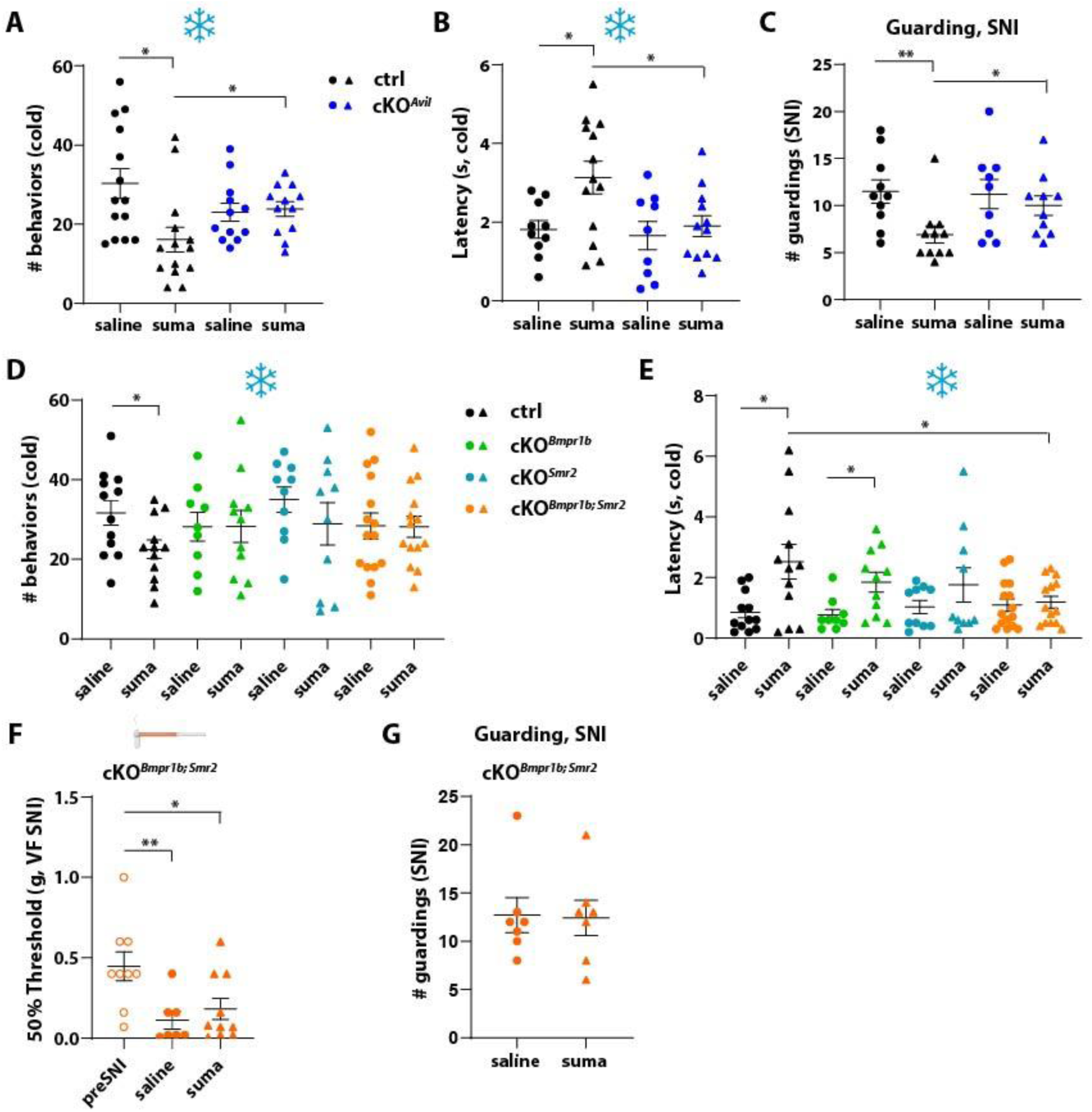
The analgesic effect of sumatriptan is mediated by Htr1b in sensory neurons, particularly the *Bmpr1b^+^* and *Smr2^+^* Aδ-HTMR subpopulations. (**A, B**) In the cold plate assay, the sumatriptan-mediated reduction of nocifensive behaviors (A) and the increase of latency (B) in control mice was abolished in the *Htr1b* cKO*^Avil^* mice. (**C**) Sumatriptan reduced guarding behaviors in control mice but not in *Htr1b* cKO *^Avil^* mice after SNI surgery. (**D, E**) In the cold plate assay, the analgesic effect of sumatriptan was partially diminished in the *Htr1b* cKO *^Bmpr1b^* or cKO *^Smr2^* mice and completely abolished in the cKO *^Bmpr1b; Smr2^* animals. (**F**) SNI-induced mechanical allodynia, reflected by reduced 50% threshold in the VF test, was not alleviated by sumatriptan in the *Htr1b* cKO *^Bmpr1b; Smr2^* animals. (**G**) Sumatriptan failed to reduce guarding behaviors in the *Htr1b* cKO *^Bmpr1b; Smr2^* animals after SNI surgery.

As *Htr1b* is broadly expressed in several transcriptionally and functionally distinct subtypes of CGRP^+^ sensory neurons (Figure 1B), we next sought to determine which of the subtypes are important for sumatriptan-mediated analgesia. Although the C-fiber CGRP^+^ neuron subtypes are traditionally considered “nociceptors”, we have found that the A fiber-HTMR subtypes (marked by the expression of Bmpr1b or Smr2) are fast conducting neurons that respond most robustly to noxious mechanical stimuli and thus may be classified as myelinated nociceptors ^18^. In an *in vivo* calcium imaging assay in which GCaMP was specifically expressed in *Bmpr1b^+^* or *Smr2^+^*DRG neurons (*Bmpr1b^Cre^; Calca^FlpE^; Ai195* or *Smr2^Cre^; Ai148*, respectively), a noxious cold stimulus (0°C) was delivered to the glabrous skin of the mouse hind paw and calcium responses of *Bmpr1b^+^*or *Smr2^+^* L4 DRG neurons were assessed. A subset of *Bmpr1b^+^*or *Smr2^+^* neurons responded to the noxious cold stimuli (Figure S6H, I), implicating these neurons as mediators of noxious cold pain. Using mice harboring *Bmpr1b^Cre^*, *Smr2^Cre^*, and *Htr1b^fl^* alleles, we generated *Bmpr1b^Cre^; Htr1b^fl/fl^* (cKO*^Bmpr1b^*), *Smr2^Cre^; Htr1b^fl/fl^* (cKO*^Smr2^*), and *Bmpr1b^Cre^*; *Smr2^Cre^*; *Htr1b^fl/fl^* (cKO*^Bmpr1b; Smr2^*) mice, which lack *Htr1b* in either *Bmpr1b^+^*, *Smr2^+^*, or both *Bmpr1b^+^* and *Smr2^+^* A fiber HTMR DRG neurons. A high degree of knock-out efficiency of Htr1b in these mutant mice was verified using RNAscope (Figure S6J, K). The cKO*^Bmpr1b^*, cKO*^Smr2^*, and cKO*^Bmpr1b; Smr2^* animals behaved grossly normal and showed comparable locomotor activity to control littermates (Figure S6L, M).

We performed the cold plate assay on the single- or double A-fiber HTMR subtype *Htr1b* cKO mice and their control littermates treated with either saline or sumatriptan. Again, sumatriptan reduced nocifensive behaviors in control mice, however it did not do so in either single- or double-subtype *Htr1b* cKO mice (Figure 4D). Sumatriptan did increase the behavioral latency in the cold plate assay in *Htr1b* cKO*^Bmpr1b^* mice (Figure 4E). In *Htr1b* cKO*^Smr2^*mice, the sumatriptan-treated group trended towards an increased latency compared to the saline treated group, although this was not statistically significant (Figure 4E). The double A-fiber HTMR-subtype cKO group (*Htr1b* cKO*^Bmpr1b; Smr2^*) exhibited insensitivity to sumatriptan, as reflected by a comparable number of behaviors and a similar latency in responses to noxious cold compared to control littermates (Figure 4D, E). Due to the similar functional properties of both subtypes, for subsequent behavioral testing, we used *Htr1b* cKO*^Bmpr1b; Smr2^* mice, which lack Htr1b in both A-fiber HTMR subtypes.

We next performed SNI surgery on cKO*^Bmpr1b; Smr2^* mice to induce neuropathic pain. Remarkably, in the double mutants, sumatriptan failed to attenuate static or punctate mechanical allodynia as well as the persistent guarding behavior (Figure 4F, G), in contrast to its analgesic effects in wildtype animals (Figure 2H, 2K, 4C). These findings indicate that Htr1b in somatosensory neurons, particularly the large diameter *Bmpr1b^+^* and *Smr2^+^* A-fiber HTMRs, is the principal target of sumatriptan for attenuating pain. This conclusion is consistent with our *in vitro* neuronal excitability measurements (Figure 1) showing that large diameter CGRP^+^ neurons are more sensitive to sumatriptan’s effects on attenuating excitability compared to small diameter CGRP^+^ DRG neurons (Figure S3A).

### Htr1b dependent and independent adverse side effects of high dose sumatriptan

Although triptans including sumatriptan have been the first line treatment for migraine, there are common adverse side effects associated with triptans including dyspnea, dizziness, paresthesia, and chest pain or tightness, which limit their clinical utility ^65–67^. We sought to test for adverse effects of sumatriptan in mice and ask whether they are mediated by Htr1b. Preclinical studies have shown that rats receiving subcutaneous sumatriptan at a dose of 12.5 mg/kg did not exhibit overt adverse effects while 25 mg/kg caused moderate behavioral depression ^68^.

When sumatriptan was given to mice at a dose of 30 mg/kg, which is 100 times greater than the dose that conferred modest analgesia (300 ug/kg), animals exhibited normal heart rate (Figure S7A), consistent with prior preclinical findings that sumatriptan is minimally vasoactive outside the trigeminal system ^68,69^. This high dose of sumatriptan reduced ambulatory activity in mice, as inferred from slightly reduced travel distance and substantially reduced rearing behaviors in the open field test (Figure 5A, B). The time animals spent in the center of the chamber, grooming or motionless was not different between saline- and high dose sumatriptan-treated groups (Figure 5C, S7B, S7C), suggesting that the high dose of sumatriptan did not cause anxiety or sedation. In a novel object exploration test, mice receiving the high dose of sumatriptan spent considerably less time exploring a novel object (Figure 5D). In the balance beam test, high dose sumatriptan-treated mice required more time to cross the beam and slipped more often while crossing (Figure 5E, F). These findings suggest that a high dose of sumatriptan caused moderately compromised well-being, manifesting in reduced ambulatory activity and rearing, reduced exploration of a novel object, and moderately impaired balance.

**Figure 5.**
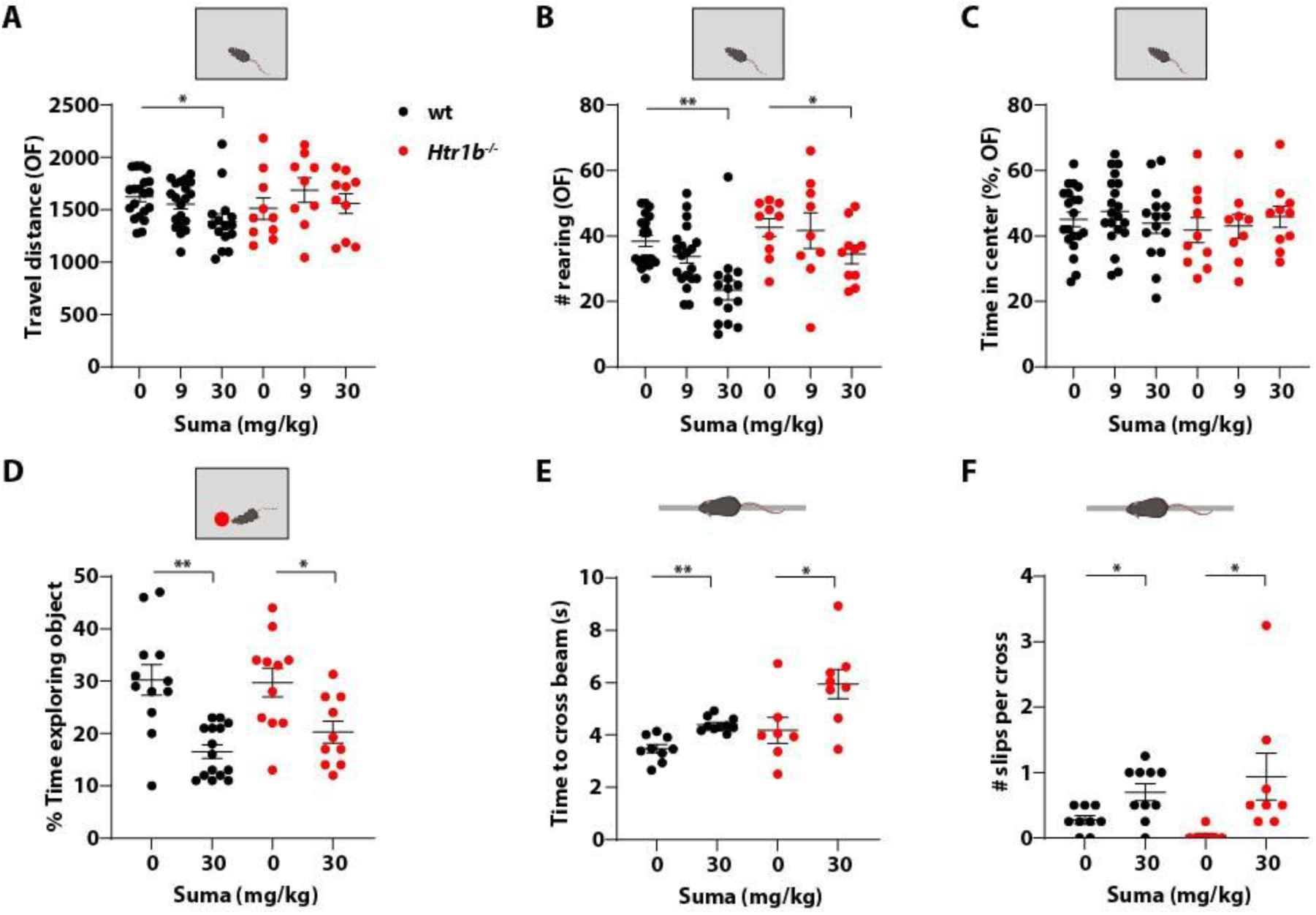
Adverse effects associated with a high dose of sumatriptan are at least partially due to Htr1b-independent off-target effects. (**A-C**) A high dose of sumatriptan (30 mg/kg) reduced ambulatory activities in the open field assay. The travel distance was slightly reduced in wildtype (wt) but not *Htr1b^-/-^* mice (A), rearing behaviors were significantly reduced in both wt and *Htr1b^-/-^* mice (B), while the time spent in the center of the chamber was not affected (C). (**D**) When given a high dose of sumatriptan, both wt and *Htr1b^-/-^*mice exhibited significantly less interest in a novel object, spending much less time exploring the object. (**E, F**) Both wt and *Htr1b^-/-^*mice exhibited impaired performance in a balance beam test when given a high dose of sumatriptan, demonstrating longer crossing time (E) and more slips during the crossing (F).

To ask whether Htr1b is responsible for these adverse effects, we generated constitutive or whole animal *Htr1b* knock-out mice (*Htr1b^-/-^*) by excising the *Htr1b* gene from the *Htr1b^flox^* allele using a germline Cre driver, EIIa-Cre ^70^. The *Htr1b^-/-^* mice lacked Htr1b expression as determined by RNAscope (Figure S7D). These animals exhibited grossly normal behavior in the open field test compared to control mice, with comparable travel distance, rearing behaviors, time in the center of chamber or motionless time (Figure 5A-C, S7C), although *Htr1b^-/-^* mice did spend more time grooming (Figure S7B).

When given a high dose of sumatriptan, *Htr1b^-/-^* mice traveled comparable distances to saline-treated *Htr1b^-/-^* mice (Figure 5A) and spent a similar amount of time in the center (Figure 5C). However, high dose sumatriptan-treated *Htr1b^-/-^* mice showed reduced rearing behaviors compared to the saline-treated *Htr1b^-/-^* controls (Figure 5B). The high dose of sumatriptan also significantly reduced exploration of a novel object in *Htr1b^-/-^* mice (Figure 5D). Moreover, sumatriptan-treated *Htr1b^-/-^* mice performed less proficiently in the balance beam test, exhibiting a longer crossing time with more slips during crossing (Figure 5E, F). We conclude that many of the adverse effects associated with a high dose of sumatriptan are due to target(s) other than Htr1b. Considering that triptans are Htr1b/1d-selective agonists, we speculate that Htr1d may be the “off-target”, although we cannot rule out the possibility of other serotonin receptors, despite having a much lower affinity, or some other unknown target, underlie these effects.

### Another triptan, zolmitriptan, acts via Htr1b in somatosensory neurons to reduce pain and exhibits Htr1b-independent adverse effects

In addition to sumatriptan, other triptans have been used to treat migraine. One of these, zolmitriptan, is more lipophilic than sumatriptan and has a longer half-life and higher oral bioavailability ^71^. Therefore, we tested zolmitriptan for its activity in a subset of behavioral assays and whether Htr1b expressed in sensory neurons is its site of action.

Using the *in vitro* calcium imaging assay, we found that zolmitriptan (10 μM) inhibited neuronal excitability of CGRP^+^ neurons to a similar extent as sumatriptan (Figure 6A, B). In addition, the cell size distribution of zolmitriptan-responding neurons was comparable to that of sumatriptan-treated neurons (Figure 6C). In behavioral assays, zolmitriptan attenuated noxious cold pain at a much lower dose (7.5-25 μg/kg) compared to sumatriptan (300 μg/kg), demonstrating its higher potency but comparable efficacy (Figure 6D). In the SNI-induced neuropathic pain model, zolmitriptan alleviated spontaneous pain as measured by a reduction in guarding behaviors (Figure 6E). Surprisingly, and different from what we had observed with sumatriptan, static mechanical allodynia was unaffected by zolmitriptan (Figure 6F). In fact, zolmitriptan slightly reduced the mechanical threshold in preSNI animals (Figure 6F), presumably due to a central action of zolmitriptan, as sumatriptan, which is more peripherally restricted, did not affect mechanical thresholds prior to surgery (Fig 2F, H). This observation is also consistent with the more severe adverse effects of zolmitriptan compared to sumatriptan (Fig 6H-M), also possibly originating from a central action. Moreover, although it is unclear how zolmitriptan caused a reduced mechanical threshold, it is possible that this effect masks any potential analgesic effect mediated by Htr1b through acting on sensory neurons. The analgesic effect of zolmitriptan in the cold plate assay was abolished by deleting *Htr1b* in both *Bmpr1b^+^* and *Smr2^+^* neurons (Figure 6G), suggesting that Htr1b expressed in A-fiber HTMR DRG sensory neurons mediates this drug’s analgesic effects.

**Figure 6.**
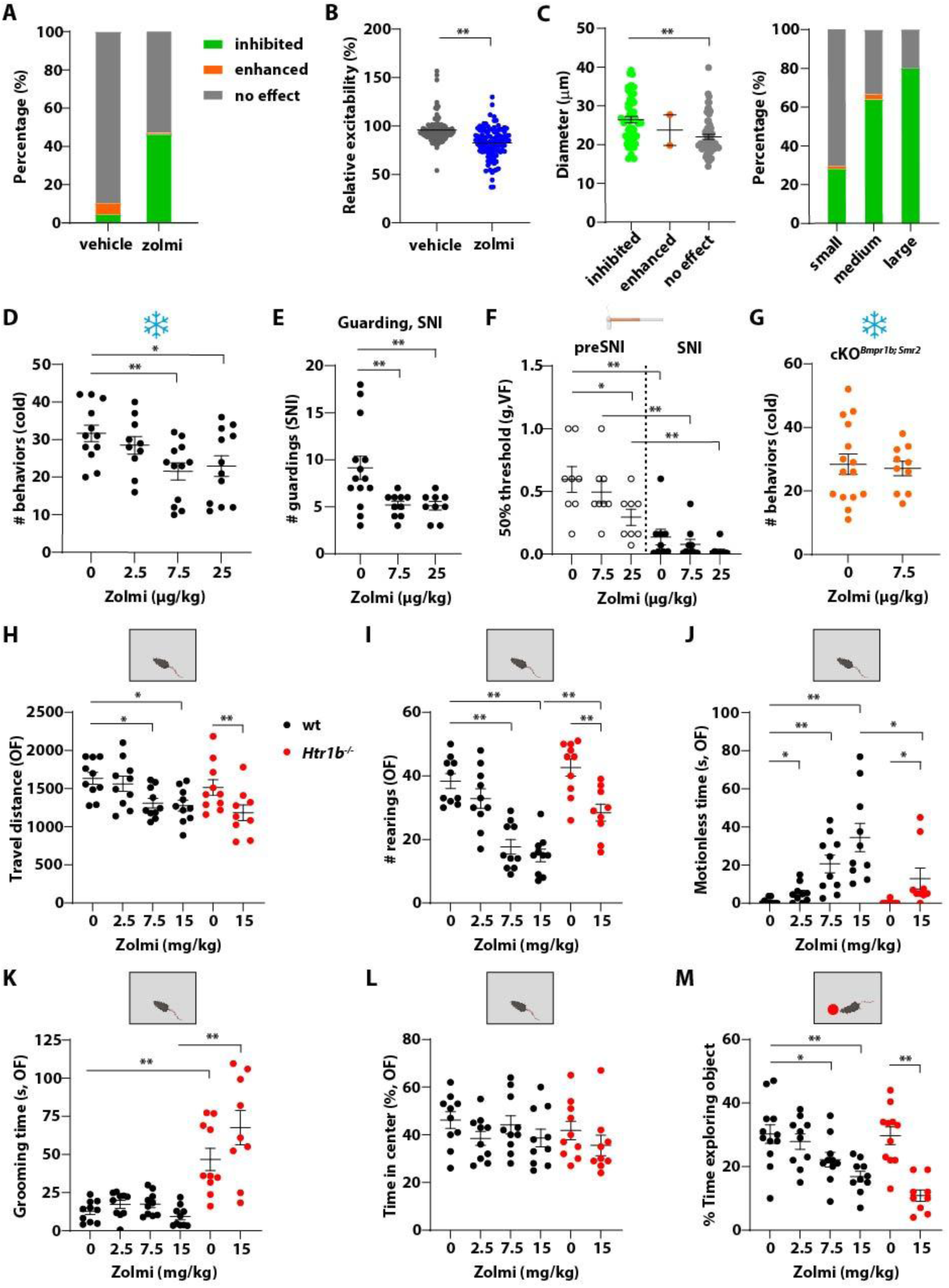
Zolmitriptan attenuates neuronal excitability and exhibits comparable analgesic effects to sumatriptan in an Htr1b-dependent manner, although it exhibits more severe adverse effects at high doses. (**A**) Over 40% of CGRP^+^ neurons were inhibited by zolmitriptan (10 μM) in the *in vitro* calcium imaging assay (N=3). (**B**) Zolmitriptan (10 μM) attenuated excitability of CGRP^+^ neurons. (**C**) Large dimeter CGRP^+^ neurons were more responsive to zolmitriptan. (**D**) Zolmitriptan dose-dependently attenuated noxious cold pain in the cold plate test. (**E**) Zolmitriptan reduced guarding behaviors in SNI mice. (**F**) Zolmitriptan did not affect mechanical allodynia after SNI surgery. (**G**) The analgesic effect of zolmitriptan in the cold plate assay was lost in the *Htr1b* cKO*^Bmpr1b; Smr2^* animals. (**H-L**) In the open field assay, zolmitriptan dose-dependently reduced ambulatory activities of both wt and *Htr1b^-/-^* mice. Travel distance (H) and rearing behaviors (I) were reduced after high dose zolmitriptan treatment. High doses of zolmitriptan also triggered a motionless state in both wt and *Htr1b^-/-^* mice that resembles sedation (J). Grooming (K) and time in the center (L) was not affected by the high doses of zolmitriptan. (**M**) High doses of zolmitriptan significantly reduced novel object exploring in both wt and *Htr1b^-/-^* mice.

When given at a dose of 7.5 mg/kg or higher, zolmitriptan reduced travel distance and rearing behaviors in the open field test (Figure 6H, I). The high dose zolmitriptan-treated animals appeared subdued and less active, displaying a longer motionless state even at the dose of 2.5 mg/kg (Figure 6J) and this effect plateaued at 15 mg/kg (Figure 6J), although grooming or time in the center was unaffected by the drug (Figure 6K, L). Zolmitriptan also dose-dependently reduced novel object exploration time (Figure 6M). Thus, overall, zolmitriptan conferred similar but more severe adverse effects compared to sumatriptan, even at low doses. Taken together, these findings indicate that zolmitriptan induces more severe adverse effects than sumatriptan, potentially due to higher blood brain barrier permeability ^71^ and access to the central nervous system (CNS).

We next tested the effects of a high dose of zolmitriptan in *Htr1b^-/-^*mice. Zolmitriptan administration to *Htr1b^-/-^* mice reduced travel distance and rearing behaviors in the open field test (Figure 6H, I). Drug-treated *Htr1b^-/-^* mice also demonstrated a longer motionless state compared to saline-treated *Htr1b^-/-^* controls, but this was less severe compared to that of wildtype mice (Figure 6J). The high dose of zolmitriptan also significantly reduced the time in which *Htr1b^-/-^* animals explored a novel object (Figure 6M). Therefore, as with sumatriptan, zolmitriptan-associated adverse effects are mostly attributed to an Htr1b-independent site of action. The finding that some zolmitriptan-triggered adverse behaviors in *Htr1b^-/-^*mice were less severe than those in wildtype mice suggests that at least some adverse effects are associated with Htr1b, possibly due to a central action of zolmitriptan.

### Gpr19, an orphan GPCR, is a promising new target for treating pain

Several orphan GPCRs emerge from our screening as interesting candidates, including Gpr19, which exhibits co-expression with CGRP in both mouse and human DRGs (Figure S1I, L) and constitutive G_i/o_-coupling activity *in vitro* (Figure S2G). Moreover, the putative Gpr19 agonist, adropin, inhibits excitability of CGRP^+^ DRG neurons *in vitro* (Figure 1F, G). To ask whether Gpr19 is the target of adropin in inhibiting CGRP^+^ neurons, we generated *Gpr19* cKO*^Calca^*mice (*Calca^CreER^; Ai148; Gpr19^fl/fl^*) and confirmed efficient depletion of *Gpr19* using *in situ* hybridization (Figure 7A, B). In the *in vitro* calcium imaging assay, the inhibitory effects of adropin on CGRP^+^ neuron excitability were diminished but not abolished in neurons obtained from *Gpr19* cKO*^Calca^* mice (Figure 7C, D), suggesting that adropin acts at least partially through Gpr19 to inhibit neuronal excitability. Although *Gpr19* is mainly expressed in C-fiber neurons in the mouse DRG (Figure S1I, left), adropin inhibited small, medium and large diameter neurons comparably (Figure S3D), further suggesting both Gpr19 and non-Gpr19 sites of adropin action.

**Figure 7.**
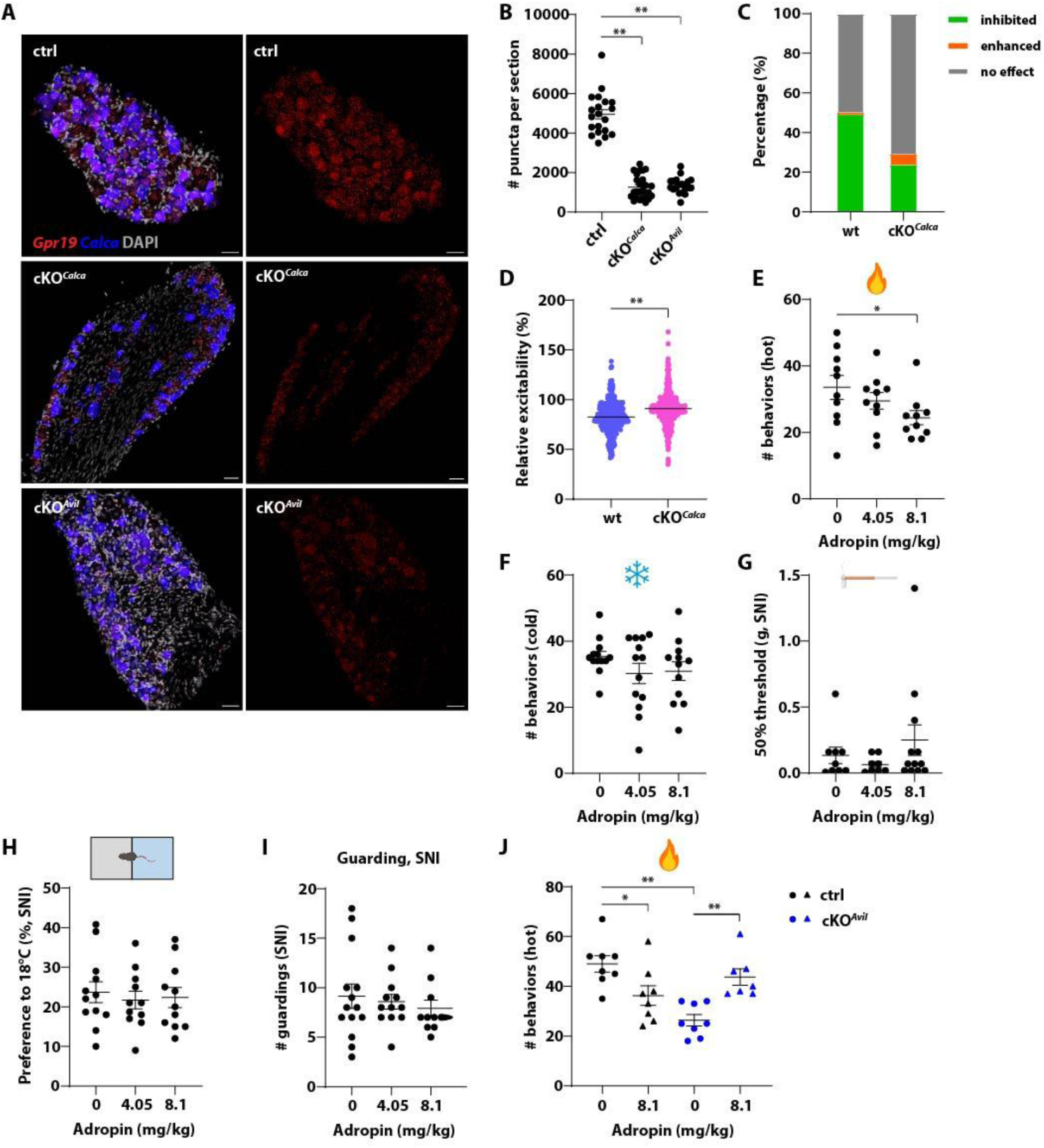
Gpr19 is a potential drug target for treating pain. (**A**) RNAscope confirmed efficient deletion of *Gpr19* in the DRG using either *Calca^CreER^* or *Avil^CreER^*. (**B**) Quantification of (A). N=3 for each genotype. (**C**) The percentage of CGRP^+^ neurons inhibited by adropin (100 nM) was reduced in *Gpr19* cKO*^Calca^*mice (*Calca^CreER^; Ai148; Gpr19^fl/fl^*) compared to control mice. (**D**) The average relative excitability of adropin-treated CGRP^+^ neurons was higher in *Gpr19* cKO*^Calca^* mice than controls. (**E**) Adropin dose-dependently reduced noxious heat-induced nocifensive behaviors. (**F**) Adropin did not affect acute noxious cold pain. (**G-I**) Adropin had no effect on mechanical allodynia as tested using VF filaments (G), cold hypersensitivity in the temperature preference assay (30°C vs 18°C, H), or spontaneous pain-like guarding behavior (I). (**J**) Adropin (8.1 mg/kg) attenuated noxious heat pain in control mice but augmented nocifensive behaviors in *Gpr19* cKO*^Avil^* mice (*Avil^CreER^; Gpr19^fl/fl^*).

*In vivo*, adropin moderately attenuated nocifensive responses to noxious heat (Figure 7E) but not noxious cold (Figure 7F), consistent with the expression of *Gpr19* in the C-Heat sensory neuron subtype marked by *Sstr2* (Figure S1I, left). In the SNI-induced neuropathic pain model, adropin had no effect on mechanical allodynia (Figure 7G), cold hypersensitivity (Figure 7H) or the spontaneous pain-like guarding behavior (Figure 7I). To determine the site-of-action of adropin, we generated *Gpr19* cKO*^Avil^*mice, in which *Gpr19* was specifically depleted in peripheral nervous system neurons including DRG neurons (*Avil^CreER^; Gpr19^fl/fl^*, Figure 7A, B). Surprisingly, the *Gpr19* cKO*^Avil^*mice exhibited hyposensitivity to heat compared to control littermates (Figure 7J). This may suggest an unknown role of Gpr19 in noxious heat detection, although the underlying mechanism is unclear. Nevertheless, in *Gpr19* cKO*^Avil^* mice, the analgesic effects of adropin in alleviating noxious heat pain were completely lost (Figure 7J). In fact, adropin augmented noxifensive responses to noxious heat in *Gpr19* cKO*^Avil^* mice (Figure 7J), suggesting that adropin may be acting on non-Gpr19 targets. This is consistent with the finding that the *in vitro* activity of adropin was only partially lost in *Gpr19* cKO*^Calca^* mice (Figure 7C, D). These findings suggest that the analgesic effect of adropin is mediated by Gpr19 in the sensory neurons, while a potential off-target(s) site of action of adropin may be pro-nocifensive.

## Discussion

We have identified a large number of GPCRs that are selectively expressed in CGRP^+^ DRG sensory neurons, and upon testing, we found that activating G_i/o_-coupled GPCRs in sensory neurons, including the serotonin receptor Htr1b, attenuates excitability of these neurons. Focusing on Htr1b, the triptans, agonists for both Htr1b and Htr1d, were found to exhibit efficacy in reducing measures of pain in a manner dependent on Htr1b expressed in CGRP^+^ sensory neurons, particularly fast-conducting A-fiber CGRP^+^ HTMRs, also known as Aδ-HTMRs or myelinated nociceptors. High doses of two distinct triptans also promoted adverse behavioral effects in uninjured mice, and these adverse effects of the triptans were largely due to an Htr1b-independent mode of action. Moreover, targeting another GPCR, Gpr19, implicates it as a potentially druggable target for treating pain. These findings establish a screening platform for identifying analgesic agonists of GPCRs expressed in DRG neurons, show that Htr1b agonists may be clinically useful beyond treating migraine, and suggest that agonists for Gpr19 and other GPCRs identified herein may be useful for treating different types of pain.

GPCRs are common drug targets. Indeed, of all clinically marketed drugs, over one-third target GPCRs ^72^. The G_i/o_-coupled GPCRs are unique because of their inhibitory mode of action. Almost all analgesic GPCR agonists activate G_i/o_-coupled receptors, including but not limited to the opioid receptors, cannabinoid receptors, α2-adrenergic receptors, and metabotropic GABA receptors ^9,10^. Besides these well characterized analgesic GPCRs, recent studies have reported other G_i/o_-coupled receptors, such as the adenosine A3 receptor (A3R) and Gpr171, and their agonists with *in vivo* analgesic efficacy ^73–76^. However, few of these studies have defined the exact site-of-action and thus the underlying mechanisms remain elusive. Instead of globally searching for G_i/o_-coupled GPCRs, we focused on the DRG sensory neurons, where nociception is initiated, and developed a pipeline for identifying druggable G_i/o_-coupled GPCRs enriched in nociceptors to treat pain. This involved bioinformatically identifying G_i/o_-coupled or orphan GPCRs selectively expressed in CGRP^+^ sensory neurons, followed by validating expression in human DRGs, evaluating the capacity of agonists/putative agonists to silence DRG neuron excitability *in vitro*, and testing compounds *in vivo* using animal behavior paradigms and conditional receptor loss-of-function alleles to determine the site of action of both desirable and undesirable behavioral consequences of the treatments. This approach is enabled by a mouse DRG neuron genetic toolkit ^5,18,20,77–81^, which enables manipulation of select sensory neuron subtypes and the roles they play when deciphering the mechanism of drug action. With the aid of these genetic tools, we can identify not only the neuronal subtype(s) responsible for the drug action but also relevant targets within the cell type. Using the serotonin receptor Htr1b and triptans as an example, conditional loss-of-function experiments revealed Htr1b expressed in CGRP^+^ sensory neurons, and in particular fast conducting A-fiber nociceptors, as the site of analgesic action of triptans. Htr1b, but not Htr1d, mediates the analgesic effects of triptans in these assays, while the adverse effects associated with high doses of triptans are mostly attributed to sites of action other than Htr1b.

Our findings revealed that triptans exhibit analgesic effects in somatic pain models in mice, including acute noxious cold pain and neuropathic pain. Triptans have been primarily used clinically for migraine treatment. In preclinical models, studies have shown varying results regarding sumatriptan’s analgesic effects in somatic or visceral pain models, indicating little to no effects on acute noxious heat and mechanical pain but efficacy in alleviating certain types of visceral pain as well as inflammatory pain ^47,82,83^. Limited studies have reported mixed results showing efficacy or lack-of-efficacy of triptans in treating neuropathic pain in preclinical models ^44–47^, possibly due to complications arising from different animal models and different methods used to induce neuropathic pain (e.g. chemotherapy vs various types of nerve injury). Our findings that triptans attenuate noxious cold pain and SNI-induced neuropathic pain are in line with the expression of *Htr1b* in the DRG across all axial level and suggest a potentially broader therapeutic utility than migraine treatment. Indeed, although triptans are the classic first line treatment for migraine, their mechanism of action and target cell type for treating migraine are unclear ^84,85^ as is their potential utility for other types of pain management. In the trigeminal system, attempts have been made to address the mechanism-of-action of the triptans. There are at least three proposed modes of action for the triptans for migraine treatment: vasoconstriction of vascular smooth muscle, inhibition of trigeminal neurons and peripheral release of neuropeptides including CGRP, and inhibition of central neurotransmitter release in the trigeminocervical complex ^62–64,86^. Related to this, CGRP inhibitors are a new class of drugs for treating migraine ^87^. Our findings that *Htr1b* is broadly expressed in CGRP^+^ sensory neurons in both mouse and human DRGs, triptans silence CGRP^+^ sensory neurons *in vitro*, sumatriptan decreases glutamate release from central sensory neuron synapses in the spinal cord dorsal horn, and Htr1b functions cell-autonomously in somatosensory neurons to mediate the analgesic properties of triptans support the view that CGRP^+^ sensory neurons are the primary targets of triptans.

One intriguing finding is the modality specific analgesic effects of Htr1b agonism, despite the broad expression of *Htr1b* in most CGRP^+^ sensory neuron subtypes (Figure 1B). Sumatriptan specifically attenuates noxious cold, but not noxious heat or acute mechanical pain (sA-E, S4A, S4B). The lack of efficacy on noxious heat and mechanical pain is consistent with previous findings ^47,83^. Our functional characterizations of DRG neurons revealed no single discrete DRG neuron subtype that exclusively encodes noxious cold ^18,20^. Rather, responses to noxious cold temperatures appear distributed across several subtypes, including subsets of Aδ-HTMRs (*Bmpr1b^+^*), Aδ-HTMR/Heat (*Smr2^+^*), C-Heat (*Sstr2^+^*), and C-HTMR/Heat (*Mrgprb4^+^*) neurons, while C-LTMRs encodes a relative change (decrease) of temperature and C-Cold thermoreceptors (*Trpm8^+^*) respond to both the decrease of temperature and innocuous to noxious cold temperatures ^18,20^. The distribution of noxious cold sensing neurons across small percentages of several subtypes and the relatively broad expression of *Htr1b* among those neurons may account for the efficacy of triptans in alleviating noxious cold pain. On the other hand, there are at least six sensory neuron subtypes that respond robustly to noxious heat ^18,20^. Among the heat-sensing neurons, only two subpopulations, the C-Heat (*Sstr2^+^*) and Aδ-HTMR/Heat (*Smr2^+^*), express *Htr1b*, while four C-fiber subtypes (the C-HTMR/Heat (*Mrgprd^+^*), C-HTMR/Heat (*Mrgprb4^+^*), C-HTMR/Heat (*Mrgpra3^+^*) and C-HTMR/Heat (*Sst^+^*) subtypes) do not (Figure 1B). Therefore, activating Htr1b in a small, select population of heat-responding neurons may not be sufficient to block heat-induced nocifensive behaviors. It is unclear why sumatriptan had no effect on noxious mechanical pain in the pinprick assay, considering our conditional gene targeting findings implicating Aδ-HTMRs in mediating the actions of sumatriptan in allodynia. Pinprick evokes a very strong reflex response, and so it is possible that moderately inhibiting the relevant sensory neuron populations may not be sufficient to block this acute, robust behavior. We did not observe any reduction of inflammatory mechanical and thermal pain in sumatriptan-treated animals (Figure 2F, G). Since sumatriptan was administered systemically through i.p. injection, our findings are consistent with prior reports implicating intrathecal but not systemic application of sumatriptan in attenuating inflammatory pain ^47^. Interestingly, compared to the acute pain models, triptans were more effective in the neuropathic pain model in alleviating static mechanical allodynia, cold hypersensitivity, and spontaneous pain (Figure 2H-M, S4E-G), despite previous findings showing that sumatriptan was ineffective in mechanical allodynia in SNI animals ^47^. It is unclear what caused this discrepancy and whether it was due to differences in performing the SNI surgery (sparing the sural vs the tibial nerve). The precise mechanism underlying the analgesic effect of sumatriptan in neuropathic pain is not clear. One caveat is that we cannot be confident about whether expression or localization of *Htr1b* is altered under neuropathic pain states. We also cannot exclude a potential contribution of changes in the central circuits after nerve injury. Future experiments are needed to address these questions.

Our findings suggest the need to re-assess the utility of existing Htr1b agonists for treating both nociceptive and neuropathic pain and that Htr1b should be considered a target for new analgesic development. In addition to Htr1b, we identified several other G_i/o_-coupled GPCRs, as well as orphan receptors that may potentially be G_i/o_-coupled, such as Gpr19. Moreover, we found that putative agonists of orphan GPCRs, including Gpr19, attenuated neuronal excitability of CGRP^+^ sensory neurons *in vitro*, a finding that merits further investigation with pain behavioral tests and conditional receptor knock out approaches. Indeed, the putative Gpr19 agonist, adropin, exhibited moderate analgesic effects in the noxious heat pain paradigm, despite potential off-target effects. Using conditional loss-of-function experiments, we found that the analgesic effects of adropin rely on Gpr19 in the sensory neurons, while off-target effects mediated by other receptors and/or excitatory signaling pathways may be pro-nocifensive. These findings point to Gpr19 as a potentially promising new target for treating pain, and other, more selective agonists of Gpr19 may prove to be therapeutically valuable. Notably, the *Gpr19* cKO mice exhibited hyposensitivity to noxious heat, suggesting that Gpr19 is involved in detection of noxious heat stimuli. Together, our findings both highlight the feasibility and reveal new opportunities for targeting GPCRs expressed in DRG sensory neurons for developing new pain therapeutics.

### Limitations of the study

Limitations of this work include the lack of understanding of the human DRG somatosensory neuron types in which the GPCRs examined are expressed; this is due to a lack of human DRG neuron subtype marker genes corresponding to physiological and functionally defined sensory neurons. In addition, our analysis was restricted to only a subset of GPCRs expressed in human DRGs, leaving other potentially relevant GPCRs unexamined. Another limitation is the absence of identified agonists for several potentially interesting orphan GPCRs. Finally, other subclasses of GPCRs expressed in DRG neurons, including G_s_- or G_q/11_-coupled receptors, may also have therapeutic potential but were not explored in this study.

## Resource availability

### Lead contact

Further information and requests for resources and reagents should be directed to and will be fulfilled by the lead contact, David Ginty.

### Material availability

The *Gpr19^fl^*mouse line is available upon request.

### Data and code availability

All data reported in this study and code used for analysis will be shared by the lead contact upon request.

## Acknowledgements

We thank Dr. Yotam Sagi (Rockefeller University) for the *Htr1b^fl^* mice, Elke Bentley from the Woolf lab (Boston Children’s Hospital) for help with conducting the Hargreaves assay, Dr. Meridith Skiba and Dr. Andrew Kruse (Harvard Medical School) for some of the GPCR-expressing plasmids, Caiying Guo at the HHMI Janelia Research Campus for generating the *Gpr19^fl^* mouse line, and Ginty lab members for discussions and comments on the manuscript. This work was supported by NIH grants NS097344-05 (D.D.G.), NS132196-02 (D.D.G.), AT011447 (D.D.G.), NS105076-01 (C.J.W), the Hock E. Tan and Lisa Yang Center for Autism Research (D.D.G.), the Edward R. and Anne G. Lefler Center for Neurodegenerative Disorders (D.D.G.), and Ono Pharmaceutical. D.D.G. is an investigator of the Howard Hughes Medical Institute (HHMI).

## Author contributions

J.P., N.S. and D.D.G. conceived the study. J.P. performed RNAscope on mouse and human DRGs, *in vitro* calcium imaging, *in vitro* cAMP signing assay and all animal behavior tests except the paw luminance and Hargreaves assay, with help from B.T.S. and M.M.D. A.M.C. performed the slice electrophysiology experiments. J.X. performed the *in vivo* calcium imaging. X.Z. performed the paw luminance assay and the Hargreaves assay under the supervision of C.J.W. L.Q. and K.L. contributed to the setup of the *in vitro* calcium imaging and the pinprick assay, respectively. J.P. and D.D.G wrote the paper with input from all authors.

## Supplemental information

Document S1. Figures S1–S7, Table S1

## Star★Methods

### Key resources table

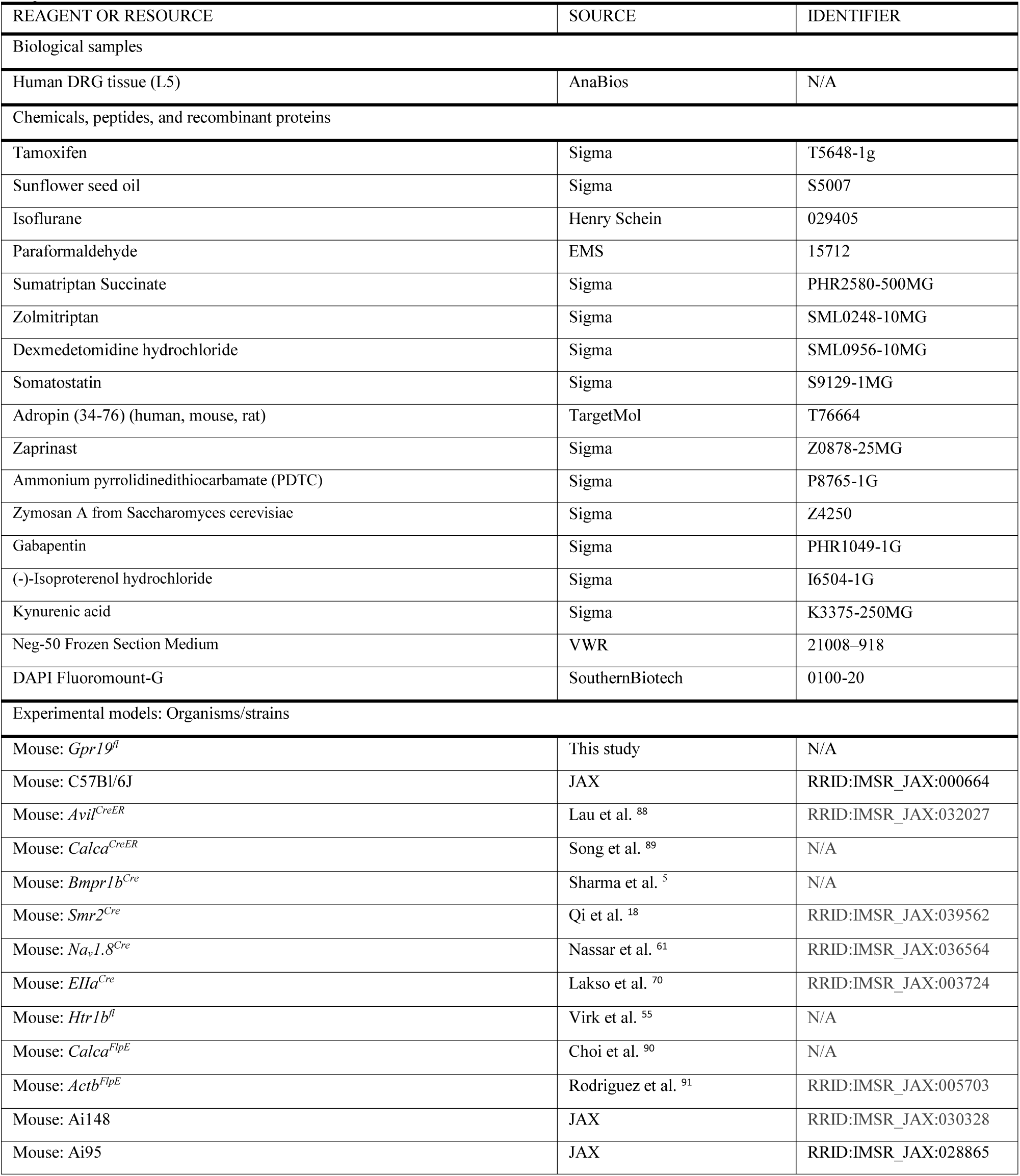
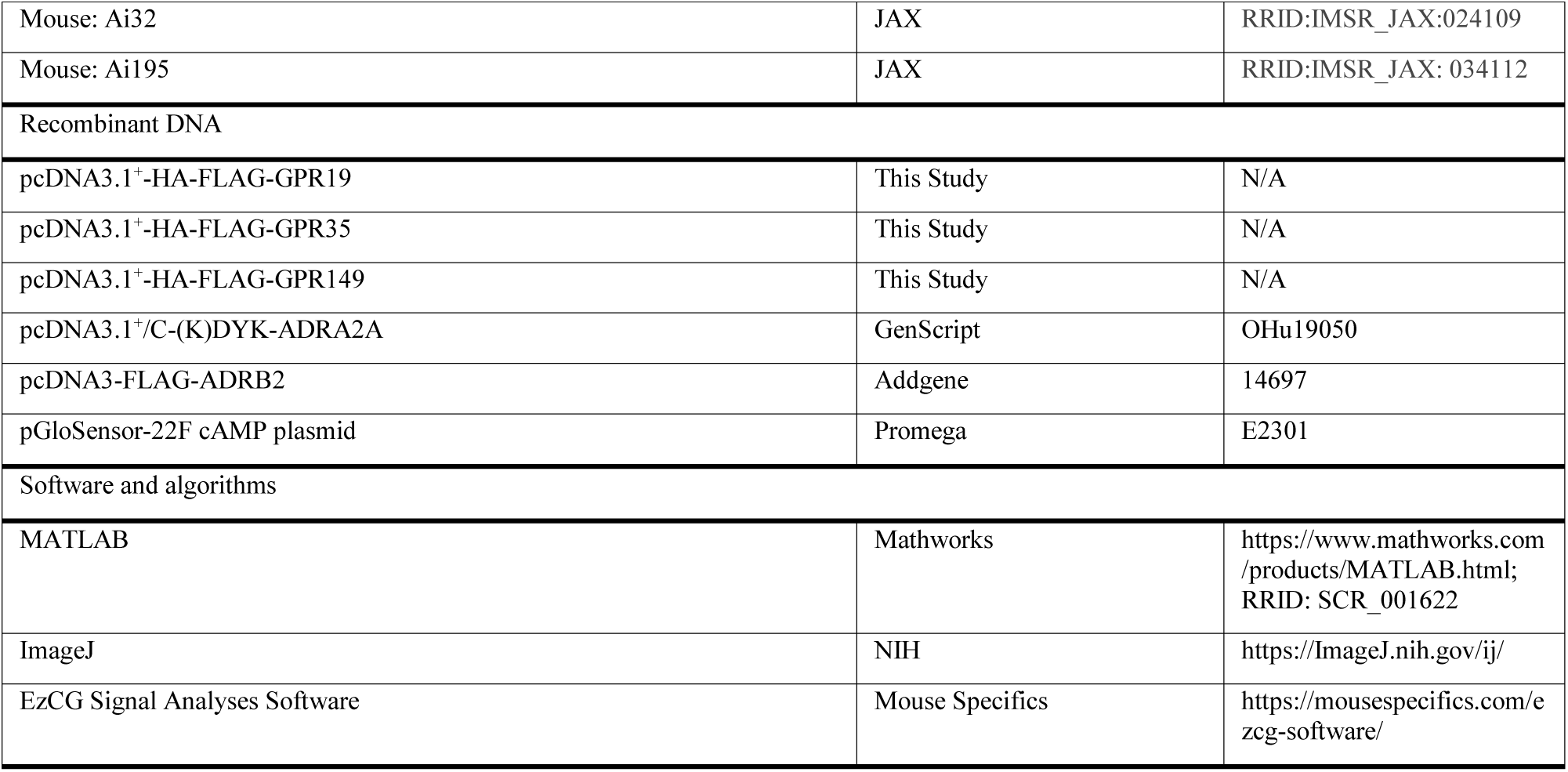

### Experimental model and subject details

Mice were used as the animal model and human DRG tissue from donors was used to assess the conservation of expression. All animal-related procedures were approved by the Harvard Medical School Institutional Animal Care and Use Committees (IACUC) and complied with the Guide for Animal Care and Use of Laboratory Animals. Both male and female animals were used for all experiments. Animals were randomly assigned to experimental groups with balanced sex representation. Only naïve adult animals were used for behavior tests and each animal received only one treatment except for the paw luminance assay where a cross-over approach was used: the mice received either saline or sumatriptan at 15 days post-SNI surgery for the paw luminance test, and two weeks later the mice received the opposite treatment at 29 days post-surgery. All SNI mice were tested by dynamic brush at 7 days post-surgery to confirm the development of allodynia. SNI mice with an allodynia score of 1.5 or higher were used for subsequent tests within at least a one-week interval, whereas mice with a score lower than 1.5 were excluded. Otherwise, all mice including outliers were included in the analyses.

The *Gpr19^fl^* mouse line was generated in this study at the Gene Targeting and Transgenics Facility at Janelia Research Campus using standard homologous recombination techniques in hybrid mouse embryonic stem (ES) cells. The only coding exon (exon 5) of *Gpr19* was flanked by loxP sites. Chimera mice were generated by blastocyst injection. Germline transmission was confirmed by genotyping and the NeoR/KanR selection cassette was removed by crossing to a germline Flp line, *Actb^FlpE^* ^91^.

## Methods details

### Human DRG tissue

The L5 human DRG tissue was purchased from AnaBios. The tissue was collected from a healthy human donor without neurological disease, and freshly frozen using dry-ice cooled isopentane. Upon arrival, the tissue was embedded in Neg-50 and stored at -80°C.

### Drug treatments

Tamoxifen solution was prepared as previously described. To generate *Htr1b* or *Gpr19* conditional knock-out mice using *Avil^CreER^*or *Calca^CreER^*, 1 mg of tamoxifen was delivered daily by intraperitoneal injection for 5 consecutive days from postnatal day 28. For calcium imaging of wildtype mice using *Calca^CreER^*, 1 mg of tamoxifen was delivered intraperitoneally one week before dissection for DRGs.

Sumatriptan succinate (Sigma) was dissolved in saline and intraperitoneally (i.p.) injected 30 min before behavior assays at 300 μg/kg for most of the behavior tests or 30 mg/kg for high-dose experiments unless otherwise specified.

Zymosan (Sigma) was dissolved in saline at 5 mg/mL. 20 μL of zymosan solution was injected subcutaneously into the plantar surface of the left hind paw.

Gabapentin was dissolved in saline and i.p. injected 30 min before behavioral test at 30 mg/kg.

### *In situ* hybridization (RNAscope)

The standard protocol from ACDBio website was used with a few modifications. Mouse DRGs were freshly dissected, embedded in Neg-50 and flash frozen. Freshly frozen mouse and human DRGs were cryosectioned at 20 μm thickness. Slides were fixed in prechilled 4% PFA at 4°C for 1 hr, and then dehydrated in 50%, 70% and 100% ethanol sequentially. Next the sections were treated with RNAscope Hydrogen Peroxide for 10 min and then digested using Protease III for 25 min. RNAscope probes were added onto the slide and incubated at 40°C for 2 hrs, followed amplification and HRP development according to the protocol. The slides were then mounted using DAPI Fluoromount-G (Southern Biotech). Imaging was done using a confocal microscope (Zeiss LSM 900), and ImageJ was used for quantification.

### *In vitro* cAMP signaling assay

The GloSensor cAMP assay (Promega) was used to assess the cAMP level in live cells. HEK293 cells were cultured using standard protocol. 1.5×10^4^ cells were seeded per well in tissue culture-treated 96-well white plates with clear flat bottom. Plates were incubated in a 37°C incubator overnight. Cells were then transfected with 50-80 ng of the pGloSensor-22F cAMP plasmid and the plasmid over-expressing human GPCRs of interest each using 500 ng of polyethylenimine per well. Cells were incubated for 24 hr. On the day of experiment, the cells were pre-equilibrated in the equilibration medium (100 μL per well, containing 88 μL CO_2_-independent media, 10 μL fetal bovine serum and 2 μL GloSensor cAMP Reagent) for 2 hr in room temperature. In a plate-reader (FlexStation3), baseline luminescence was acquired before adding compounds. Cells were pre-treated with varying concentrations of agonists by adding 1 μL of 100× compound stock solution per well with 3 replicate wells. After 5 min incubation, 1 μL of 100× isoproterenol stock solution (10 μM) was added to all wells. Luminescence was measured at 5 min to 20 min post-isoproterenol addition. For ligand-independent constitutive activity assessment, baseline luminescence was acquired before compound treatment, and luminescence measurements were taken at multiple time points from 5-20 min after isoproterenol addition with 6 replicate wells. Each experiment was repeated at least twice.

### DRG neuron dissociation and culture

One week after tamoxifen treatment or two weeks from the first tamoxifen treatment for the *Htr1b* conditional knock-out mice, animals were sacrificed and DRGs were immediately dissected in cold HBSS buffer. DRGs were collected in 1 mL of digestion solution (HBSS buffer containing 5 mg/mL dispase, 2 mg/mL collagenase and 0.1 mg/mL DNase I) on ice, and then digested by incubating at 37°C under constant rotation (60 rpm) for 45 min. Digestants were spun down at 200 rcf for 5 min and washed twice with HBSS. Cells were resuspended in 1 mL of DMEM medium (Gibco) containing 1/100 Penicillin-Streptomycin and 1/10 fetal bovine serum, and triturated using fire polished glass pipettes from large to small diameters sequentially. 100 µL of dissociated cell suspension was plated onto each 12 mm glass coverslip that had been treated with 50 µg/ml poly D-lysine and 1 µg/ml laminin overnight in a 24-well plate. Cells were left at room temperature for 30 min to attach to the coverslip. 500 µL of plating medium (DMEM medium containing 1/100 Penicillin-Streptomycin, 1/10 fetal bovine serum, 100 µg/mL NGF, 100 µg/mL BDNF, 10 µg/mL GDNF) was added to each well. The plate was then kept in a 37°C incubator with 10% CO_2_.

### *In vitro* calcium imaging

The *in vitro* calcium imaging protocol was adapted from a previous report ^42^. After culturing for 48 hrs, plating medium was removed from the 24-well plate and fresh DMEM medium was added. After 20 min in room temperature, the medium was removed and observation solution (145 mM NaCl, 5 mM KCl, 2 mM CaCl_2_, 1 mM MgCl_2_, 10 mM HEPES pH7.0, 1/1000 Penicillin-Streptomycin) was added to the well. The coverslip was then transferred to a customized recording chamber (Warner Instrument) that was situated on an upright fluorescence microscope (Nikon) and connected to a gravity fed perfusion system with a six-channel perfusion valve controller and a pump. Cells were constantly perfused with observation solution.

The perfusion valve controller was connected to and automatically controlled by an Arduino board. After a 15 s baseline of observation solution perfusion, cells were treated with 25 mM KCl for 15 s, followed by 285 s of wash-out using observation solution. Then a second KCl treatment was delivered for 15 s, followed by 45 s of wash-out and subsequent 5 min of vehicle or drug treatment. The third KCl treatment was delivered along with vehicle or drug. Another two KCl treatments were given at a 5 min interval. Imaging was done using a camera (Thorlabs) connected to the fluorescence scope. Images were analyzed and quantified using ImageJ. Cell excitability was defined as the maximum ΔF/F value during each trial. The first two KCl trials were used to determine the baseline excitability before drug treatment. Only cells with comparable excitability for the first two trials were included in the analysis (i.e. difference of excitability < 15%). Cells whose excitability of the third trial (drug-treated trial) is 15% lower or higher than the baseline (mean of the first two trials) were considered inhibited or enhanced by the drug.

### *In vivo* calcium imaging

The *in vivo* calcium imaging experiments were performed as previously described ^18^. Mice were anesthetized with 2% inhalational isoflurane, with body temperature maintained at 37°C throughout the experiment. The back hair near the lumbar region was removed by shaving and applying Nair. After skin was cleaned and disinfected, an incision was made on top of the lumbar vertebrae, and the paravertebral muscles on L3-L5 were removed. A spinal clamp was used to stabilize the spine. The bone on top of the L4 DRG was removed with a rongeur to expose the DRG. The surgical preparation was then transferred to the platform under an upright epifluorescence microscope (Zeiss) with a 10× objective. A 470 nm LED (Thorlabs) was used as the light source. A CMOS Camera (Thorlabs) was triggered at 10 fps with 50 ms exposure time. All stimuli were synchronized with the camera and LED using a DAQ board (National Instrument). A Peltier device (13×12×2.5 mm, TE technology) was used to deliver thermal stimulation. Thermal paste (Thermal Grizzly) was evenly applied on the Peltier surface. The Peltier was then gently pressed on the glabrous skin of the hind paw to form a firm contact without applying excessive pressure. A thermocouple microprobe (Physitemp) was inserted between the Peltier and skin as a separate measurement of applied temperature. After 20 s of baseline at 32°C, temperature dropped to 0°C at the rate of 5°C s^−1^. The temperature was maintained at 0°C for 23.6 seconds before raising back to the baseline. Pinch stimuli were applied at the end of the trial to confirm the viability of cells. Only cells that responded to pinch were included in the analyses.

ImageJ plugin “moco” ^92^ and “Unsharp mask” filter were used for motion correction and spatial high-pass filtering. Individual cells were manually circled and aligned across videos for pinch and thermal stimulation. The calcium signal measurements from ImageJ were analyzed using custom MATLAB codes. Heatmap plots were generated with ΔF/F of individual cells during thermal stimulation period.

### Spared nerve injury (SNI)

The surgery was done as previously described ^48^. Mice were anesthetized using isoflurane. Eye lubricant was applied to both eyes. The hair on the left thigh was removed using Nair. The exposed skin was disinfected by 70% ethanol and then betadine. An incision was made above the knee joint. Muscles were carefully separated to expose the sciatic nerve bundle. The branching point where the sural nerve was separated from the tibial and the common peroneal was located. The two nerves were tied using non-absorbable silk suture (Henry Schein) down the branching point without touching the sural. A second knot was tied slightly lower from the first one with the absorbable vicryl suture (Covetrus), and a transection was made between the two knots. The muscles were pulled together and the skin was stitched by making 3-4 double knots with the absorbable vicryl suture. Liquid bandage was applied to the skin. The mice were placed in the recovering cage above a heating pad and put back to the home cage after they awoke. The mice were monitored for 5 days post-surgery. Allodynia in SNI mice were validated using the dynamic brush assay 7 days post-surgery. Animals with allodynia scores under 1.5 were excluded from subsequent tests.

### Open field

The open field assay was done as previously described ^93^. Two days prior to the experiment, mice were moved to the testing room to acclimate to the room and the testing chamber (25×25×18 cm, length × width × height). On the day of test, mice were singly placed in the chamber and recorded for 5 min using a top-positioned camera at 30 fps. A custom MATLAB code was used to track distance travelled and time spent exploring in the center.

### Cold and hot plate

The cold and hot plate assays were done as previously described ^90^. Animals were habituated for 20 min in a chamber (14×14×30 cm) with a metal plate floor set at 26°C for 2 days. On the day of the experiment, mice received a single dose of either vehicle or sumatriptan (30-300 μg/kg) through intraperitoneal injection. 30 min after the injection, mice were singly placed in a chamber with a clear acrylic front side. The floor metal plate was set at 0°C for cold plate or 51°C for hot plate. Mice were kept in the chamber for 1 min for cold plate or 30 s for hot plate. Behaviors were recorded using a front-positioned high-speed camera at 120 fps. Number of nocifensive behaviors (clenching, flicking, guarding or licking of fore paws and hind paws, jumping) and the latency to the first behavior were manually scored by an experimenter blinded to genotypes or treatments.

### von Frey (VF)

The von Frey assay was done as previously described ^90^. Mice were habituated for 30 min on an elevated metal grid in a clear acrylic chamber for 2 days. Punctate stimulations were delivered to the pads (for non-surgery mice) or the lateral side (for sham and SNI surgery mice) of the left hind paw using a series of filaments (0.008-4 g). For each filament, 5 stimuli were delivered, followed by another 5 after a 30 s interval. Sudden withdraw, guarding, flicking or licking of the paw indicated a positive response. The number of responses were recorded. The lowest force that elicited 5 or more responses out of 10 was considered the 50% threshold.

### Dynamic brush

Dynamic brush was done as previously described ^94^. Mice were habituated for 30 min on a metal grid in a clear acrylic chamber for 2 days. Three gentle brushes were delivered to the lateral part of the left hind paw using a paintbrush. Brushing continued with 5-10 s intervals for 3 min per trial. 3 trials were performed for each animal. The responses were graded with a score from 0 to 3: 0 for no response, 1 for sustained lifting or withdraw of the ipsilateral hind paw, 2 for lateral kicking, and 3 for liking. The maximal response for each trial was considered the score for that trial, and the scores for all three trials were averaged. The number of behaviors were also recorded. Animals with a score of 1.5 and higher were considered allodynic.

### Pinprick

The pinprick assay was done as previously described ^20^. Mice were habituated in VF chambers for 2 days before the experiment. A custom-made pinprick apparatus with a sharp pin (FST) was used. Upward movement of the pin was triggered with a manual push-button. An accelerometer (SparkFun) was attached to the wire grid floor and a circuit was set up to measure conductance between the pin and the mouse paw that rested on a conductive metal grid. A DAQ board (National Instruments) was used to synchronously record the accelerometer signals, the button presses and skin contact signals. Data were collected for 3 minutes during which an experimenter delivered multiple pinprick stimuli to the plantar surface of the hind paws. Animal behavior was also recorded with a bottom-viewed camera at 200 fps.

Custom MATLAB code was used for data analyses. Each push of a button was considered a trial. Only trials where the pin made strong contact with the paw skin (conductance value for pin-to-skin contact value exceeding 1.5 V) after the button push were included in the data analyses. Stimulus onset was defined as first contact of pin to skin. Movement onset was empirically defined as when the accelerometer value reached 0.15 V. Movement magnitude was computed as area under the curve for accelerometer values across 0.4 s after stimulus onset.

### Inflammatory pain (zymosan model)

Inflammatory pain was induced by intraplantar injection of zymosan ^95^. Zymosan solution (100 μg in 20 μL) or vehicle (saline) was subcutaneously injected into the left hind paw. Mice then received i.p. injection of saline or sumatriptan 3.5 hr post zymosan/vehicle intraplantar injection. Behavior tests were performed 30 min afterwards. VF assay was used to test mechanical pain.

The Hargreaves assay was done for thermal hyperalgesia as previously described ^96^. Animals were habituated to the Hargreaves apparatus (Ugo Basile) consisting of a glass floor with Plexiglas enclosures for 5 consecutive days (1-2 hr per day). Baseline withdrawal latencies were measured prior to treatment, after which mice were assigned to four groups using stratified allocation to ensure comparable baseline distributions and balanced sex representation. On the day of testing, mice received intraplantar injections of either saline or zymosan. At 3.5 hr post-injection, sumatriptan or saline was i.p. administered. Thermal withdrawal latency was assessed 30 min post i.p. injection (i.e. approximately 4 hr post intraplantar injection). A focused radiant heat source (50% infrared intensity) was applied to the plantar surface of the ipsilateral hind paw, and latency to withdrawal was recorded. The mean of five trials was used for latency. All behavioral testing was performed by an experimenter blinded to treatment conditions.

### Temperature preference

The temperature preference assay was done as previously described ^93^. Animals were habituated for 20 min on a metal plate floor set at 26°C for 2 days. Two metal plates (14×14×18 cm) were placed adjacent to each other. The temperature of each plate was controlled separately. A divider was placed in the middle of the chamber. Both plates were set at 30°C to test for place preference. For the cold temperature preference, one plate was set at 30°C (neutral side) and the other at 18°C (cool side). Mice were placed in the chamber and free to explore. The movement was recorded by a top camera at 30 fps and tracked using a custom MATLAB code. The time the mice spent on each side was quantified, and the preference of the cool side was plotted.

### Guarding behavior

Naïve mice were habituated in the same way as the cold/hot plate test. Guarding behaviors were assessed in SNI mice at 2-3 weeks post-surgery. 30 min post i.p. injection, mice were placed in a chamber with the metal plate floor set at 26°C. Behaviors were recorded for 5 min with a front high-speed camera at 120 fps and manually scored by an experimenter blinded to treatments. Lifting and then prolonged holding of the ipsilateral hind paw was considered guarding behavior. Care was taken to differentiate a normal step during walking from a guarding behavior. All manual scoring was done by the same blinded experimenter using the same criteria throughout the experiments.

### Paw luminance assay

The paw luminance assay was done as previously described ^51^. Mice were first habituated individually for 30 minutes in a black recording chamber (see below), followed by a 30-minute video recording session (pre-injury session), one day prior to undergoing the SNI procedure. One week after injury, mechanical allodynia was confirmed in all mice using von Frey testing. At two weeks post-injury, mice were randomly assigned to receive either vehicle or drug treatment, with balanced sex distribution across groups. Mice in the drug group received a single intraperitoneal injection of sumatriptan (300 μg/kg), while control mice received an equivalent volume of saline. Following treatment, mice were placed in the recording chamber and recorded for 30 minutes. Two weeks later, the treatments were crossed over between groups (i.e., vehicle-treated mice received sumatriptan and vice versa), followed by the same recording procedures.

The recording setup and analysis pipeline were identical to those previously described ^51^. Briefly, mice were placed in a black acrylic enclosure (18×18×15 cm) that was enclosed on all sides except the bottom. The enclosure was positioned on a 25×25×0.5 cm glass platform. Two independent near-infrared (NIR) LED strips (850 nm; Huake Light Electronics) were mounted 10 cm beneath the glass floor and powered by a 12 V DC supply. Video recordings were acquired in darkness using an NIR-sensitive camera (Basler) positioned 30 cm below the glass surface. Video acquisition was performed using Pylon Viewer software (Basler) under standardized camera settings. All recordings were conducted at 25 frames per second with an initial resolution of 1000×1000 pixels, which was subsequently downscaled to 500×500 pixels for high-throughput analysis. Paw surface contact was quantified based on luminance signals from both hind paws, and the luminance ratio was calculated as the average luminance of the injured (ipsilateral) hind paw relative to the uninjured (contralateral) hind paw over the entire 30-minute recording session (i.e., 45,000 frames).

### Texture preference (sandpaper)

The texture preference assay was done as previously described ^93^. Mice were habituated in the chamber for 2 days. Two types of sandpapers (3M) with smooth (400-grit, blue) or rough texture (100-grit, purple) were used. The chamber was divided into two compartments using a divider. Each compartment had one of the two sandpapers or construction papers with matching colors on the floor. Mice were first run in the chamber with color-matching construction papers (blue on one side and purple on the other) on the floor for 10 min (non-texture control, NT). Then the construction papers were replaced with sandpapers of the matching color. Mice were then run in the chamber for 10 min on the textured floors. Video recording was done using a top camera at 30 fps. Tracking and analysis was done using MATLAB.

### Novel object test

Mice were habituated in a chamber for 2 days before the experiment. On the day of experiment, mice were allowed to explore in the empty chamber for 5 min and put back to the home cage. A novel object (a wooden block in a novel shape) was introduced to the chamber. The mice were placed back into the chamber with the novel object and stayed for 5 min. Video recording was done using a top camera at 30 fps. The amount of time the mice spent exploring the object was manually scored by an experimenter blinded to the genotype or the treatment.

### Balance beam

The balance beam assay was done as previously described ^93^. The balance beam rod was constructed of 1 m of matte white acrylic with a flat 12 mm-wide surface. The beam rod was mounted 50 cm above the bench top. A black matte acrylic box was placed at each end of the beam to provide a start and end enclosure for the animal. A soft pad was stretched below the beam to serve as a soft-landing surface to cushion falls. The day prior to the test, animals were room habituated and trained to cross the beam freely. Each animal was placed in the center of the 12 mm-wide rod and encouraged to walk toward the end enclosure containing home cage bedding. The animals were encouraged to walk across the beam at least 3 crossings before returning to the home cage. Once trained, each animal was tested for 4 consecutive crossings with a 5 s rest between each crossing. Both top and side cameras were used to record the crossings. Time to cross and number of slips were manually scored in a blinded manner.

### Electrocardiogram

Heart rate was measured using ECGenie Clinic (Mouse Specifics). Mice were habituated on the electrode tower for 30 min. After vehicle or drug administration, mice were immediately placed on the electrode tower. Recording started 5 min post injection, and multiple readings were taken at different time points from 5-30 min post injection. Data was analyzed using the EzCG Signal Analyses software (Mouse Specifics).

### Slice electrophysiology

P15-P21 acute spinal cord slices were used for whole-cell patch clamp recordings of lamina II dorsal horn neurons as previously described ^97^. Mice were briefly anesthetized with isoflurane, and intracardially perfused with ice-cold oxygenated choline solution before spinal cord removal. The isolated spinal cord was embedded in low-melting agarose (Sigma Aldrich), and 300 μm transverse slices were prepared from lumbar levels using a Leica vibrating blade microtome (Leica). Spinal cord slices were prepared in ice-cold oxygenated choline solution containing: 92 mM choline chloride, 2.5 mM KCl, 1.2 mM NaH_2_PO_4_, 30 mM NaHCO_3_, 20 mM HEPES, 2.5 mM glucose, 5 mM sodium ascorbate, 2 mM thiourea, 3 mM sodium pyruvate, 10 mM MgSO_4_, 0.5 mM CaCl_2_. Slices recovered at 34°C for 30 min in HEPES holding solution equilibrated with 95% O_2_, 5% CO_2_, containing: 86 mM NaCl, 2.5 mM KCl, 1.2 mM NaH_2_PO_4_, 35 mM NaHCO_3_, 20 mM HEPES, 25 mM glucose, 5 mM sodium ascorbate, 2 mM thiourea, 3 mM sodium pyruvate, 1 mM MgSO_4_, 2 mM CaCl_2_ (pH 7.3, osmolarity 305) ^97^. Spinal cord slices were then transferred to a submerged recording chamber at room temperature and continuously perfused with artificial cerebrospinal fluid (aCSF) containing: 2.5 mM CaCl_2_, 1 mM NaH_2_PO_4_, 119 mM NaCl, 2.5 mM KCl, 1.3 mM MgSO4, 26 mM NaHCO_3_, 25 mM dextrose, and 1.3 mM sodium ascorbate, saturated with 95% O_2_, 5% CO_2_ at a flow rate of ∼1-2 mL/min. Patch pipettes (5–7 MΩ) were pulled from borosilicate glass (Sutter Instruments) and filled with an internal solution containing: 135 mM CsMeSO_3_, 4 mM ATP-Mg^2+^, 0.3 mM GTP-Na^+^, 1 mM EGTA, 3.3 mM QX-314(Cl^-^ salt), 8 mM Na_2_-phoshocreatine and 10 mM HEPES. Neurons were visualized using infrared differential interference contrast and fluorescence microscopy. Whole cell voltage-clamp recordings were obtained from lamina I-II_o_ dorsal horn neurons under visual guidance using a 40× water-immersion objective. Neurons were voltage-clamped at -70 mV, and input resistance and series resistance were monitored throughout each experiment. Cells were excluded from analysis if these values changed by >20% during the recording.

Excitatory post-synaptic currents (EPSCs) were evoked optogenetically by stimulating *Na_v_1.8^ChR2^* primary afferent axonal terminals with 1 ms pulses of blue light (473 nm, 5 mW) delivered at 30 s intervals using LED whole field illumination through the water-immersion 40x objective. After a 10 min stable baseline, sumatriptan (10 μM) was bath-applied for 10 min. In a subset of experiments, optically evoked EPSCs (oEPSCs) were blocked by NBQX (10 μM) confirming that they were mediated by AMPARs. Data were acquired using a Multiclamp 700B amplifier, a Digidata 1440A acquisition system, and Clampfit software (Molecular Devices). Sampling rate was 10 kHz, and data were low-pass filtered at 3 kHz. No correction for junction potential was applied. oEPSC peak amplitudes were measured, and responses were normalized to the mean of the 10 min baseline period. Normalized values were then used for average response plots, with all cells time-aligned at the beginning of the 10-min baseline. Averaged oEPSC amplitudes for the 2 min period immediately before sumatriptan application were compared with averaged EPSC amplitudes during the 2 min period at 8-10 min after sumatriptan bath-application (just before wash-out).

### Quantification and statistical analysis

Data analysis and quantification were done by manual scoring in a blinded fashion or automatic tracking using MATLAB. GraphPad Prism 10 was used for statistical analyses. All data are shown as the mean ± standard error of the mean (SEM). Comparisons between two groups were done by a two-tailed Student’s t test and significance was assigned at p < 0.05 (* for p < 0.05, ** for p <0.01).

