## Supplemental Information for "Targeting G_i/o_-coupled GPCRs to inhibit nociceptors: insights from the serotonin receptor Htr1b and triptans"

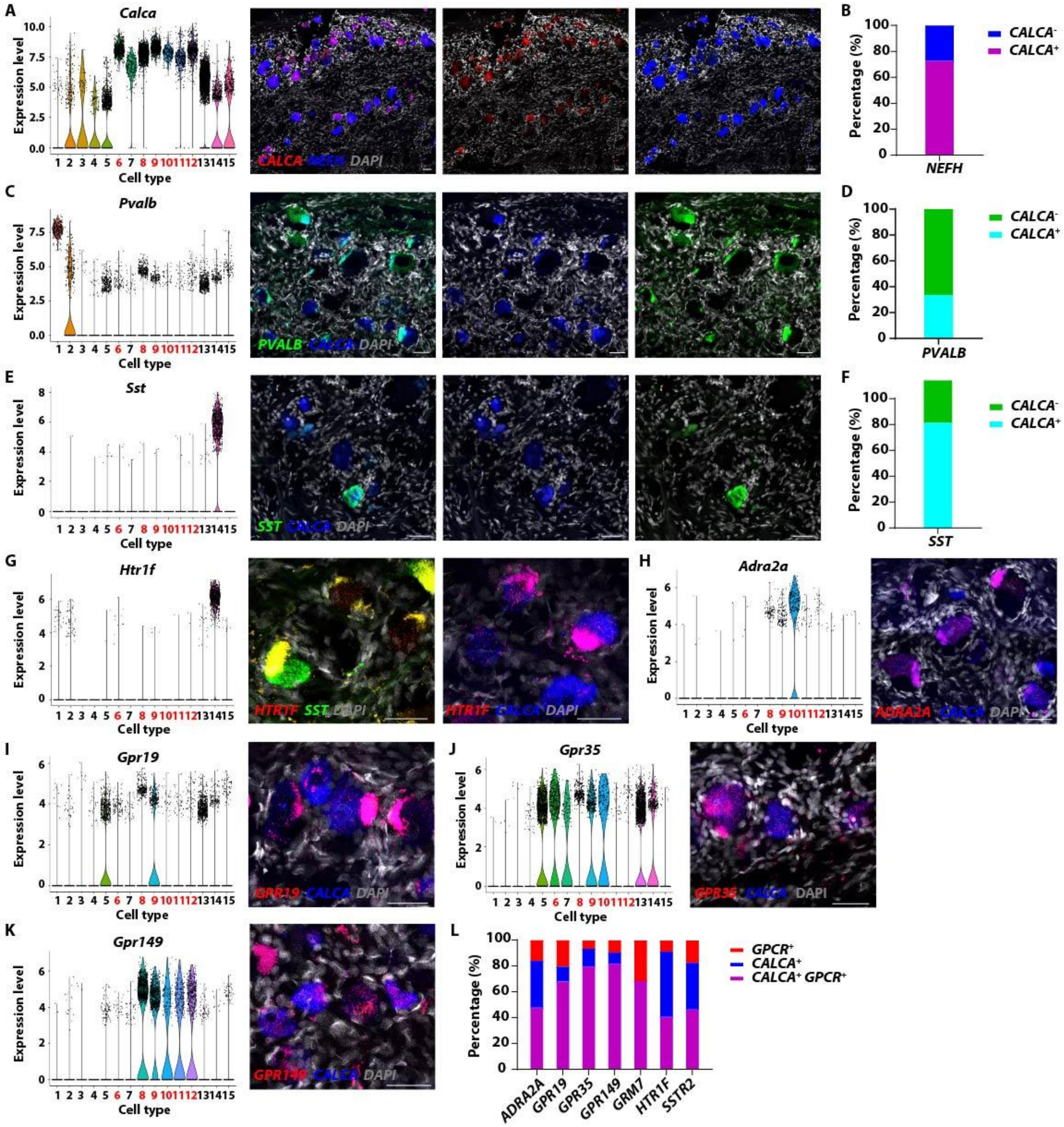


**Figure S1. Expression of a selected subset of GPCR candidates validated by RNAscope in human DRGs, related to Figure 1.** (**A-F**) Expression of the marker genes *CALCA* (A), *PVALB* (C) and *SST* (E) in human DRGs was assessed by RNAscope (right), with the expression of the mouse counterpart across all DRG neuron subtypes in our RNA-seq dataset shown for comparison (left). For all the violin plots in this figure, the cell types are number coded with the CGRP$^{+}$ subtypes highlighted in red: 1. Proprioceptors; 2. Aβ-SA LTMRs; 3. Aβ-RA LTMRs; 4.

Aδ-LTMRs; 5. C-LTMRs; 6. C-HTMR/Heat (*Mrgpra3*$^+$); 7. C-HTMR/Heat (*Mrgprb4*$^+$); 8. C-Heat (*Sstr2*$^+$); 9. CGRP-ε (*Oprk1*$^+$); 10. CGRP-γ (*Adra2a*$^+$); 11. Aδ-HTMRs (*Bmpr1b*$^+$); 12. Aδ-HTMR/Heat (*Smr2*$^+$); 13. C-HTMR/Heat (*Mrgprd*$^+$); 14. C-HTMR/Heat (*Sst*$^+$); 15. C-Cold (*Trpm8*$^+$). Quantifications were shown in (B, D, F) respectively. Scale bar: 50 μm. (**G-K**) A subset of GPCRs were co-stained with *CALCA* in human DRGs by RNAscope. Scale bar: 50 μm. (**L**) Quantification of the co-expression of GPCRs with *CALCA* in human DRGs.

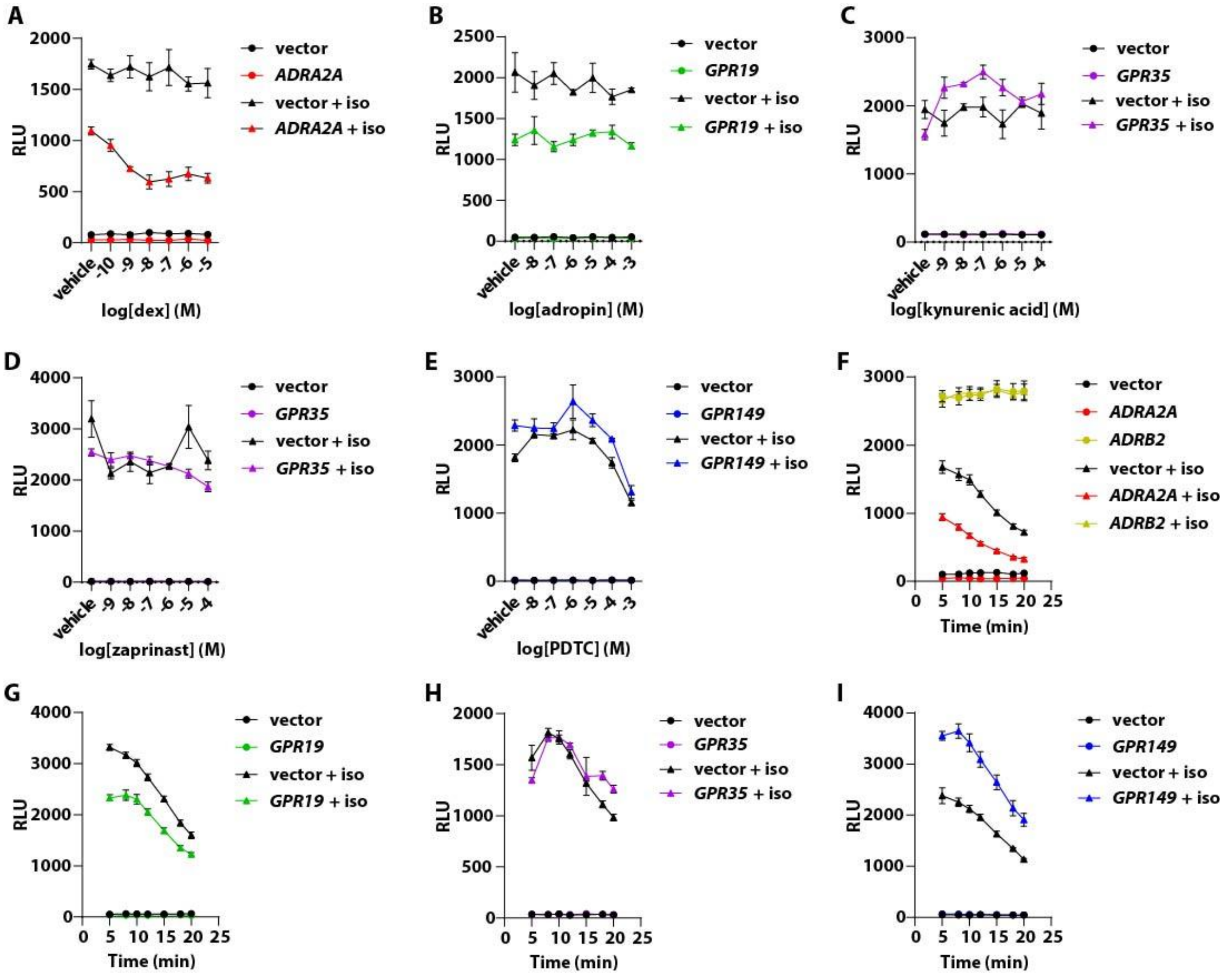


**Figure S2. Select orphan GPCRs were tested for G$_{i/o}$-coupling activities using an *in vitro* cAMP signaling assay, related to Figure 1.** (**A**) Dexmedetomidine dose-dependently inhibited cAMP accumulation in *ADRA2A* over-expressing cells. (**B**) Adropin did not affect cAMP levels in *GPR19* over-expressing cells. (**C, D**) Neither kynurenic acid (C) nor zaprinast (D) affected cAMP levels in *GPR35* over-expressing cells. (**E**) PDTC reduced cAMP accumulation at higher doses in both *GPR149* over-expressing and vector-expressing cells. (**F**) Compared to the vector-expressing control, cells over-expressing *ADRB2* had a significantly higher cAMP level, which was not further increased by its agonist isoproterenol (iso), while cells over-expressing *ADRA2A* have a lower cAMP level, consistent with their G$_s$- and G$_{i/o}$-coupling activity respectively. (**G**) Cells over-expressing *GPR19* exhibited a diminished level of cAMP, suggesting constitutive G$_{i/o}$-coupling activity. (**H**) Over-expression of *GPR35* did not affect cAMP levels. (**I**) Over-expressing *GPR149* increased cAMP levels, suggesting constitutive G$_s$-coupling activity.

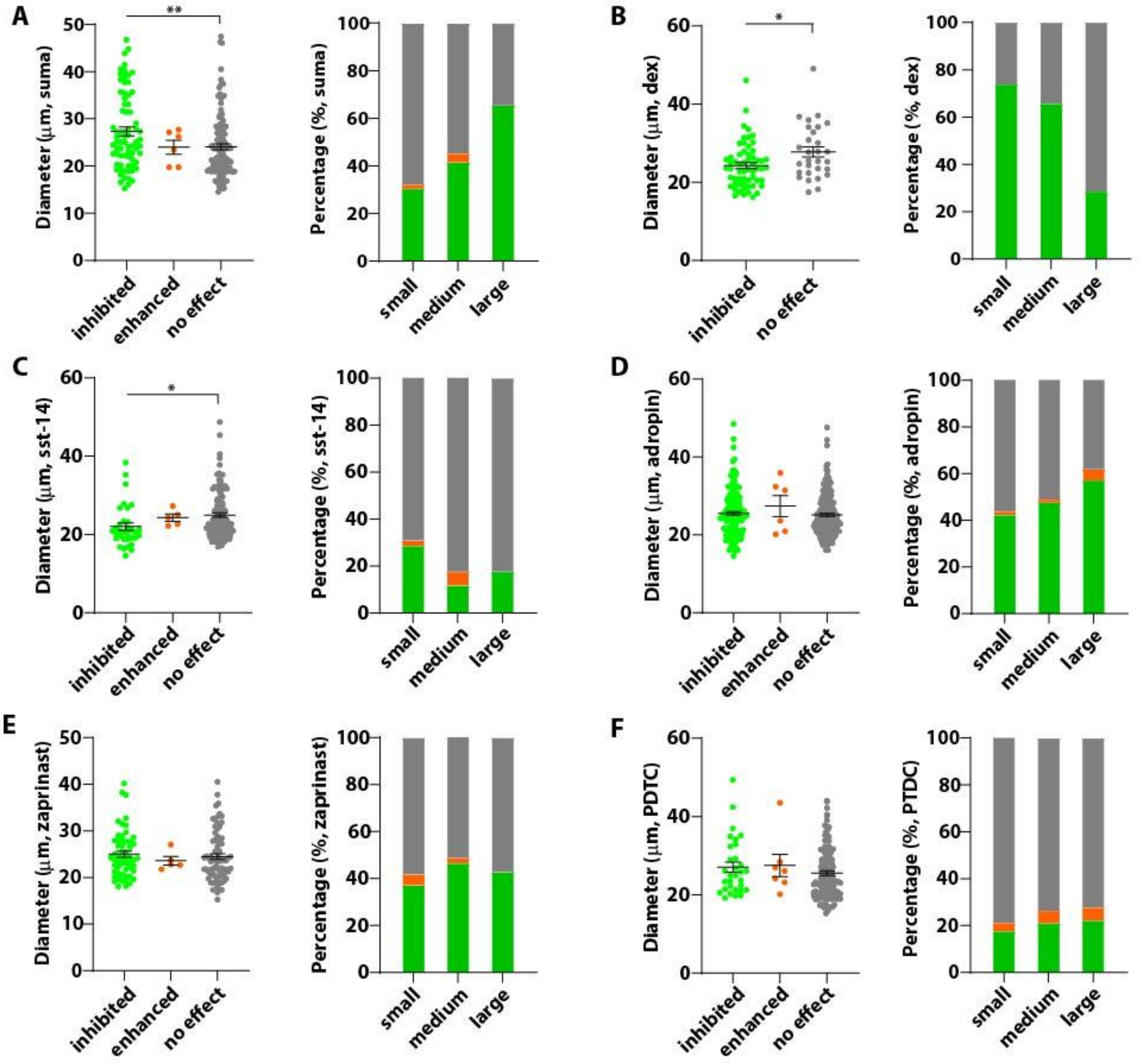


**Figure S3. Cell size distribution of the *in vitro* calcium imaging assay, related to Figure 1.** (**A**) Cells that are inhibited by sumatriptan have larger diameters than the non-responding cells (left). Each dot represents a cell. A higher percentage of large diameter cells are inhibited by sumatriptan than small- or medium-diameter cells (right). (**B, C**) Dexmedetomidine (dex, **B**) and sst-14 (**C**) preferentially inhibit small diameter cells, consistent with the expression of the α2-adrenergic receptor and the somatostatin receptor in small diameter DRG neurons. Cells treated with adropin (**D**), zaprinast (**E**) or PDTC (**F**) had a relatively uniform size distribution, indicating likely more than one targets. Green: inhibited. Orange: increased. Gray: no effect.

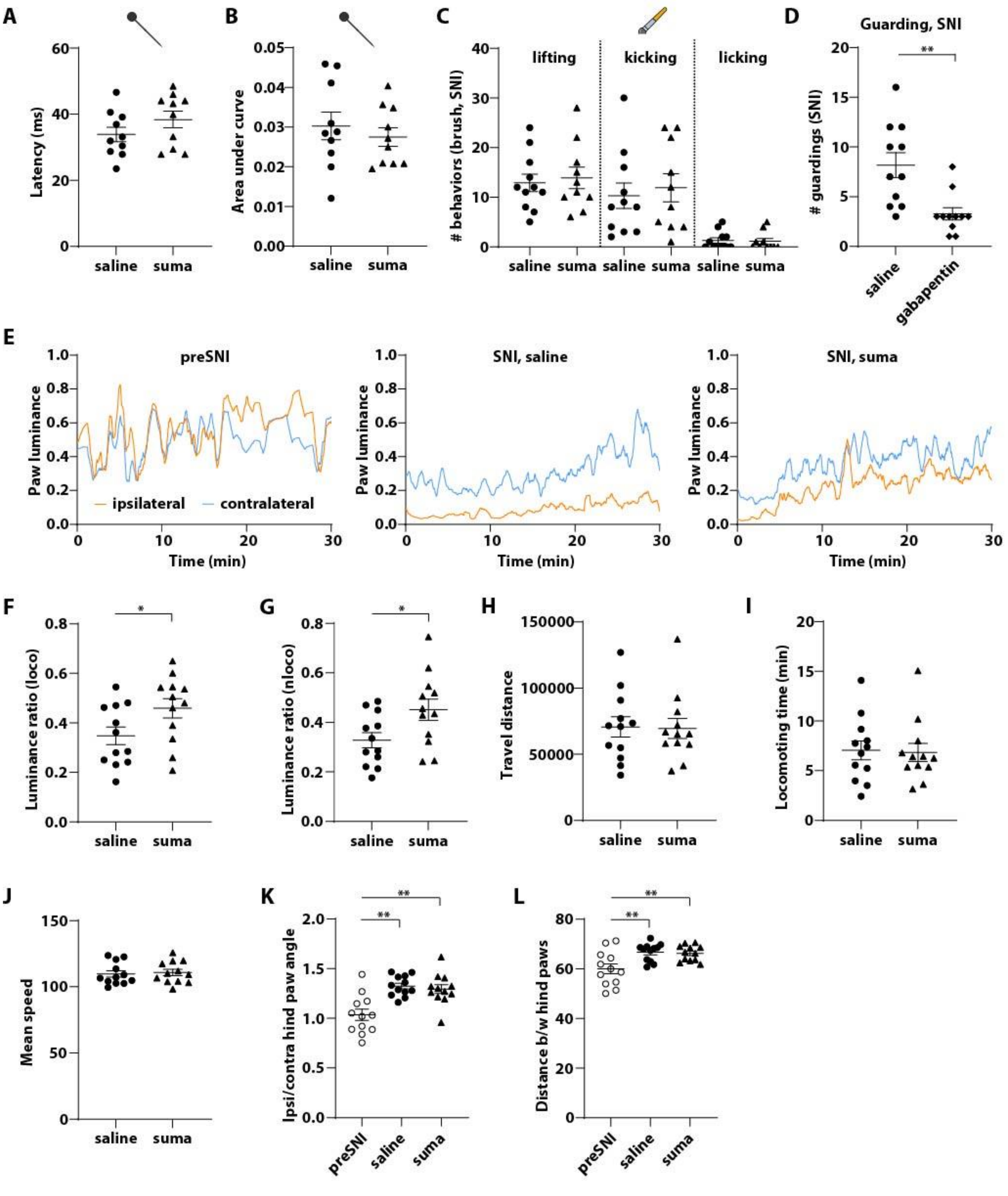


**Figure S4. Sumatriptan did not affect acute mechanical pain, dynamic allodynia, locomotion or gait of the SNI mice, related to Figure 2.** (**A, B**) Sumatriptan had no effect on acute mechanical pain in the pinprick assay. Latency (A) and cumulative response magnitude (area under the curve of response magnitude between 0-0.4 s) (B) were not different between the treatment groups. (**C**) Sumatriptan did not reduce the number of nocifensive behaviors in the dynamic brush test (**D**) Gabapentin (30 mg/kg, i.p.) significantly reduced guarding behaviors in SNI mice. (**E**) Representative traces of paw luminance of ipsilateral (orange) and contralateral hind paw (blue) before SNI (left), after SNI treated with saline (middle) or sumatriptan (right). The paw luminance values were scaled (normalized by the maximum value) and smoothed before plotting. The SNI animals had significantly reduced paw luminance of the ipsilateral paw compared to the contralateral hind paw. Sumatriptan partially alleviated the decrease of luminance of the ipsilateral hind paw. (**F, G**) Sumatriptan increased the paw luminance ratio of the SNI mice during both the locomoting (F) and non-locomoting phase (G). (**H-J**) Sumatriptan did not affect travel distance (H), locomoting time (I) or speed (J) of the SNI mice. (**K, L**) Sumatriptan did not affect the gait of the SNI mice, including the ipsilateral hind paw angle (normalized to contralateral hind paw) (K) and the distance between the hind paws (L).

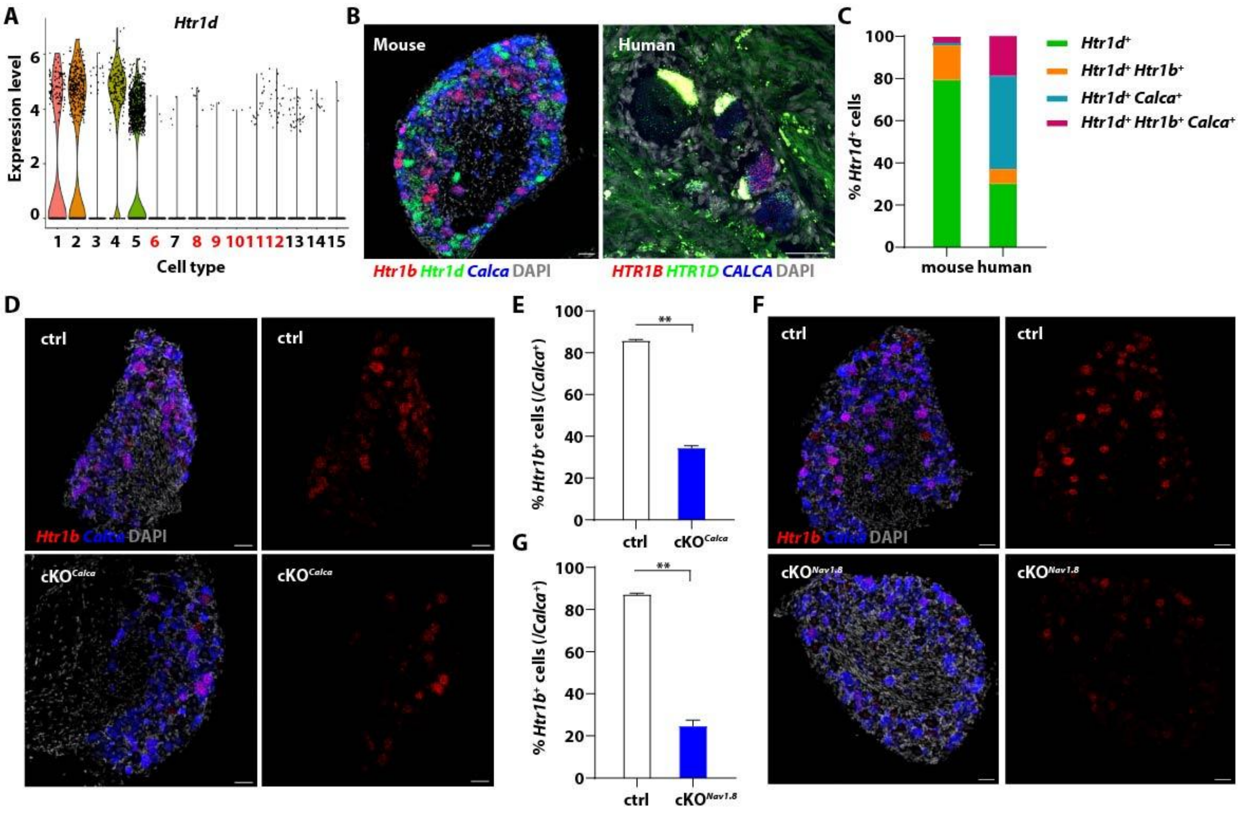


**Figure S5. Htr1b was efficiently depleted in the cKO$^{Calca}$ mice used for the *in vitro* calcium imaging and slice electrophysiology experiments, related to Figure 3.** (**A**) Expression of *Htr1d* in our mouse DRG single cell RNA-seq dataset. The cell types are number coded with the CGRP$^{+}$ subtypes highlighted in red: 1. Proprioceptors; 2. Aβ-SA LTMRs; 3. Aβ-RA LTMRs; 4. Aδ-LTMRs; 5. C-LTMRs; 6. C-HTMR/Heat (*Mrgpra3*$^{+}$); 7. C-HTMR/Heat (*Mrgprb4*$^{+}$); 8. C-Heat (*Sstr2*$^{+}$); 9. CGRP-ε (*Oprk1*$^{+}$); 10. CGRP-γ (*Adra2a*$^{+}$); 11. Aδ-HTMRs (*Bmpr1b*$^{+}$); 12. Aδ-HTMR/Heat (*Smr2*$^{+}$); 13. C-HTMR/Heat (*Mrgprd*$^{+}$); 14. C-HTMR/Heat (*Sst*$^{+}$); 15. C-Cold (*Trpm8*$^{+}$). (**B**) Expression of *Htr1b*, *Htr1d* and *Calca* was assessed by RNAscope. *Htr1d* expression did not overlap with *Htr1b* or *Calca* in mouse (left) and human DRGs (right). The arrow indicates a cell expressing *HTR1D* but not *HTR1B* or *CALCA* in the human DRG (right). Scale bar: 50 μm. (**C**) Quantification of (B, N=3 for mouse DRG and N=1 for human DRG). (**D, E**) RNAscope assay confirmed efficient depletion of *Htr1b* in the cKO$^{Calca}$ mice (N=3 for each genotype). Scale bar: 50 μm. (**F**) RNAscope showed significantly reduced *Htr1b* signal in the DRGs of the cKO$^{Nav1.8}$ mice. Scale bar: 50 μm. (**G**) Quantification of (F, N=3 for each genotype).

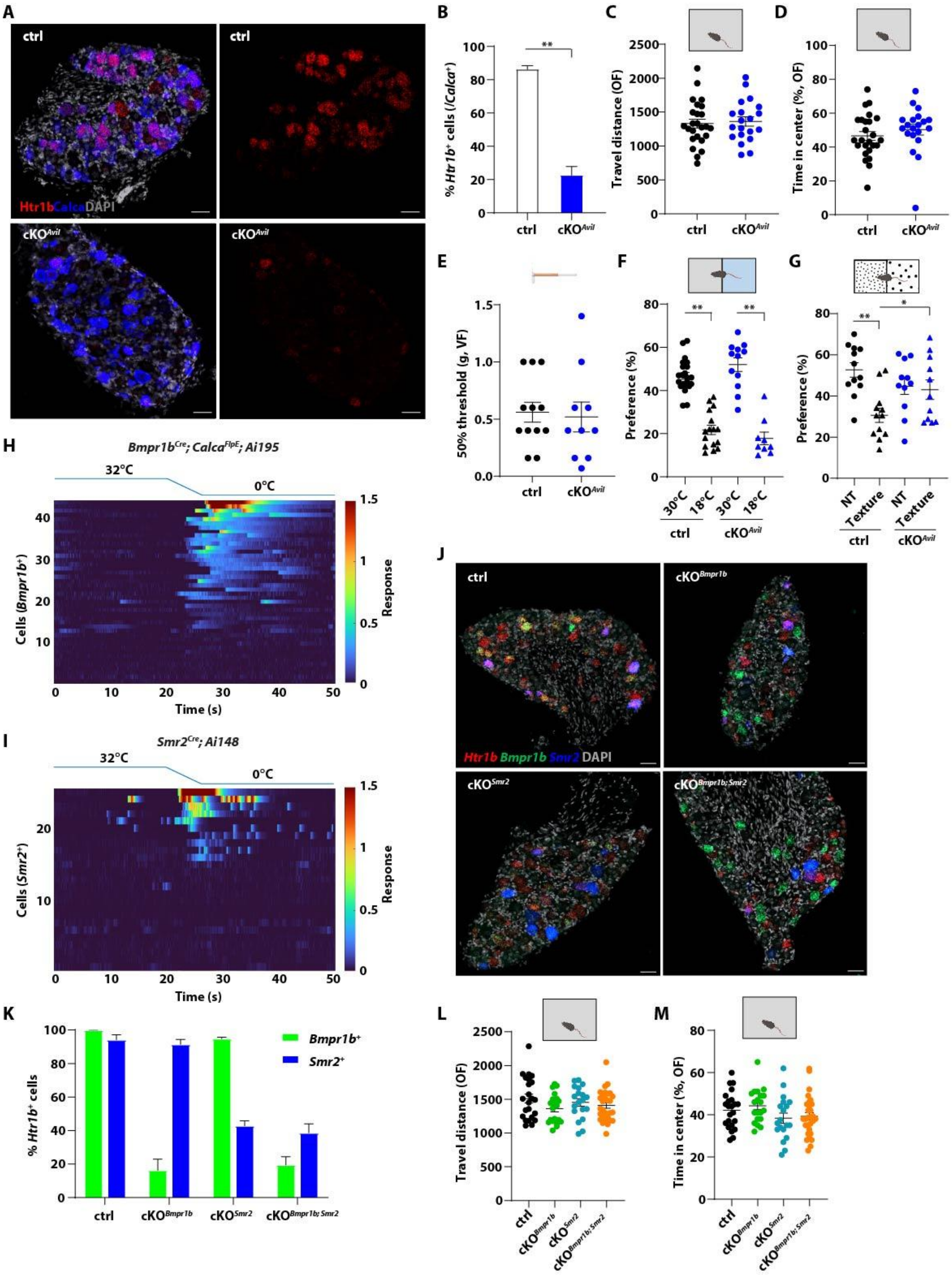


**Figure S6. Characterization of the pan-sensory neuron and subtype-specific *Htr1b* cKO mice, related to Figure 4.** (**A, B**) RNAscope confirmed efficient depletion of *Htr1b* in the cKO$^{Avil}$ mice (A), quantified in (B, N=3 for both control and cKO$^{Avil}$). Scale bar: 50 μm. (**C, D**) The *Htr1b* cKO$^{Avil}$ mice behaved normally in the open field (OF) test, exhibiting comparable travel distance (C) and time in center (D) to the control littermates. (**E, F**) The *Htr1b* cKO$^{Avil}$ mice showed normal mechanical and thermal sensitivity as tested by von Frey assay (E) and temperature preference assay (F). (**G**) In the sandpaper test, *Htr1b* cKO$^{Avil}$ mice exhibited no preference for the smooth or rough texture, in contrast to control littermates which demonstrated strong aversion to the rough texture. NT: non-texture, construction paper on both sides. Texture: sandpaper 400 vs 100 grit. Preference is defined as the percentage of time the animal spent on the rough side, i.e. 100 grit sandpaper (Texture), or the same side but with construction paper (NT). (**H, I**) A subset of *Bmpr1b*$^+$ (E, N=2, n=44) and *Smr2*$^+$ (F, N=3, n=25) DRG neurons responded to noxious cold (0°C) in the *in vivo* calcium imaging assay. Thermal stimuli were delivered to the glabrous skin of the mouse hind paw using a Peltier, while the ipsilateral L4 DRG was imaged. The thermal stimuli started with physiological skin surface temperature (32°C) as the baseline, and then ramped (5°C $s^{-1}$) and held (20 s) at 0°C. Each row of the heatmap represents responses of an individual neuron. (**J, K**) RNAscope confirmed efficient deletion of *Htr1b* in the cKO$^{Bmpr1b}$, cKO$^{Smr2}$ or cKO$^{Bmpr1b; Smr2}$ mice (J), with quantification shown in (K, N=3 for each genotype). Scale bar: 50 μm. (**L, M**) The subtype-specific *Htr1b* cKO mice performed normally in the open field test, showing comparable travel distance (L) and time in center (M) to the control littermates.

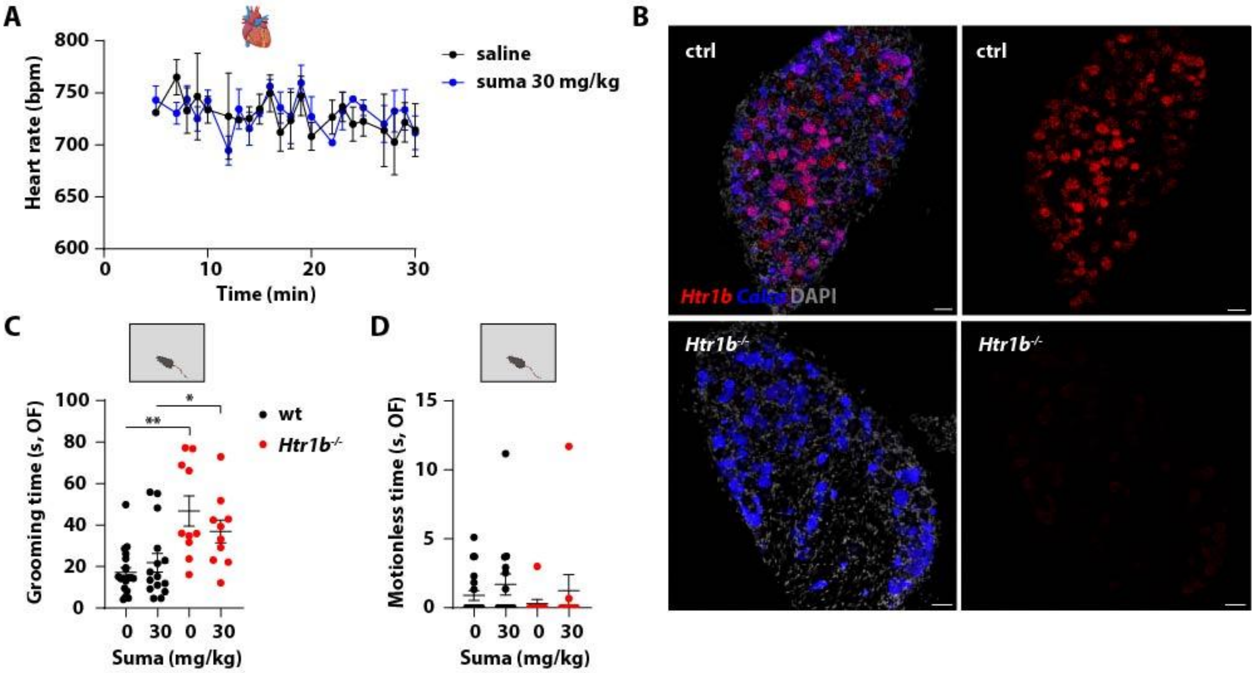


**Figure S7. A high dose of sumatriptan didn't affect cardiac function or cause sedation-like behavior, related to Figure 5.** (**A**) Mice receiving a high dose of sumatriptan (30 mg/kg) showed normal heart rate compared to saline-treated animals (N=10 for both groups). (**B**) The high dose of sumatriptan did not affect grooming behavior, although *Htr1b*$^{-/-}$ mice (red) exhibited enhanced grooming compared to the control mice (black). (**C**) The high dose of sumatriptan did not cause a significant increase of motionless time in both control (black) and *Htr1b*$^{-/-}$ mice (red). (**D**) RNAscope confirmed complete loss of *Htr1b* signal in DRGs of *Htr1b*$^{-/-}$ mice (N=3). Scale bar: 50 µm.

| Gene | Family | Expression pattern in mouse DRG | G protein coupling | Expression in human DRG |
|---|---|---|---|---|
| *Adgrf5* | Adhesion Class GPCRs | C-HTMR/Heat (*Mrgpra3*$^+$, *Mrgprd*$^+$) | $G_{q/11}$ | – |
| *Adora2b* | Adenosine receptors | C-HTMR/Heat (*Mrgpra3*$^+$, *Mrgprd*$^+$) | $G_s$, secondary: $G_{q/11}$ | – |
| *Adra2a* | Adrenoceptors | CGRP-γ (*Adra2a*$^+$), C-Heat (*Sstr2*$^+$), CGRP-ε (*Oprk1*$^+$) | $G_{i/o}$, secondary: $G_s$ | Co-expressing CGRP |
| *Agtr1b* | Angiotensin receptors | C-HTMR/Heat (*Mrgpra3*$^+$, *Sst*$^+$), CGRP-ε (*Oprk1*$^+$), CGRP-γ (*Adra2a*$^+$) | $G_{i/o}$, $G_{q/11}$ | – |
| *Avpr1a* | Vasopressin and oxytocin receptors | C-Heat (*Sstr2*$^+$) | $G_{q/11}$ | – |
| *Calcrl* | Calcitonin receptors | CGRP-ε (*Oprk1*$^+$), C-Heat (*Sstr2*$^+$), CGRP-γ (*Adra2a*$^+$), C-HTMR/Heat (*Sst*$^+$) | – | – |
| *Ccr10* | Chemokine receptors | C-Heat (*Sstr2*$^+$), C-Cold (*Trpm8*$^+$) | $G_{i/o}$ | – |
| *Crhr2* | Corticotropin-releasing factor receptors | C-Heat (*Sstr2*$^+$), C-Cold (*Trpm8*$^+$) | $G_s$, secondary: $G_{q/11}$ | – |
| *Cysltr2* | Leukotriene receptors | C-HTMR/Heat (*Sst*$^+$) | $G_{q/11}$, secondary: $G_{i/o}$ | – |
| *Drd1* | Dopamine receptors | Aδ-HTMRs (*Bmpr1b*$^+$) | $G_s$ | – |
| *F2r* | Proteinase-activated receptors | CGRP-γ (*Adra2a*$^+$) | – | – |
| *F2rl1* | Proteinase-activated receptors | C-HTMR/Heat (*Sst*$^+$) | – | – |
| *Fzd4* | Class Frizzled GPCRs | Aδ-HTMRs (*Bmpr1b*$^+$), CGRP-γ (*Adra2a*$^+$), C-HTMR/Heat (*Mrgpra3*$^+$) | $G_{12/13}$ | – |
| *Fzd5* | Class Frizzled GPCRs | C-HTMR/Heat (*Mrgpra3*$^+$, *Sst*$^+$, *Mrgprd*$^+$), CGRP-γ (*Adra2a*$^+$), CGRP-ε (*Oprk1*$^+$), C-Heat (*Sstr2*$^+$) | $G_{q/11}$ | – |
| *Galr1* | Galanin receptors | C-Heat (*Sstr2*$^+$), CGRP-ε (*Oprk1*$^+$), CGRP-γ (*Adra2a*$^+$), C-HTMR/Heat (*Mrgpra3*$^+$) | $G_{i/o}$ | Not expressed |
| *Ghsr* | Ghrelin receptor | CGRP-γ (*Adra2a*$^+$) | $G_{q/11}$, secondary: $G_{i/o}$, $G_{12/13}$ | – |
| *Gper1* | G protein-coupled estrogen receptor | C-HTMR/Heat (*Sst*$^+$) | $G_{i/o}$, secondary: $G_s$ | – |
| *Gpr135* | Class A Orphans | C-Heat (*Sstr2*$^+$), CGRP-ε (*Oprk1*$^+$), C-Cold (*Trpm8*$^+$), C-HTMR/Heat (*Sst*$^+$, *Mrgpra3*$^+$), CGRP-γ (*Adra2a*$^+$) | – | – |
| *Gpr149* | Class A Orphans | C-Heat (*Sstr2*$^+$), CGRP-ε (*Oprk1*$^+$), CGRP-γ (*Adra2a*$^+$), Aδ-HTMRs (*Bmpr1b*$^+$), Aδ-HTMR/Heat (*Smr2*$^+$) | – | Co-expressing CGRP |

| | | | | |
|---|---|---|---|---|
| *Gpr150* | Class A Orphans | C-HTMR/Heat (*Sst*$^+$, *Mrgpra3*$^+$), CGRP-γ (*Adra2a*$^+$), CGRP-ε (*Oprk1*$^+$), C-Cold (*Trpm8*$^+$) | – | – |
| *Gpr160* | Class A Orphans | CGRP-ε (*Oprk1*$^+$), C-Heat (*Sstr2*$^+$), C-HTMR/Heat (*Sst*$^+$, *Mrgprd*$^+$) | – | – |
| *Gpr165*[a] | – | C-Heat (*Sstr2*$^+$) | – | – |
| *Gpr174* | Class A Orphans | Aδ-HTMRs (*Bmpr1b*$^+$), CGRP-γ (*Adra2a*$^+$), CGRP-ε (*Oprk1*$^+$), C-Cold (*Trpm8*$^+$) | – | – |
| *Gpr179* | Class C Orphans | C-HTMR/Heat (*Mrgpra3*$^+$, *Sst*$^+$, *Mrgprd*$^+$), CGRP-ε (*Oprk1*$^+$), CGRP-γ (*Adra2a*$^+$) | – | – |
| *Gpr19* | Class A Orphans | C-Heat (*Sstr2*$^+$), C-Cold (*Trpm8*$^+$), CGRP-ε (*Oprk1*$^+$), C-HTMR/Heat (*Sst*$^+$, *Mrgpra3*$^+$), CGRP-γ (*Adra2a*$^+$) | – | Co-expressing CGRP |
| *Gpr26* | Class A Orphans | C-Heat (*Sstr2*$^+$), C-Cold (*Trpm8*$^+$) | $G_s$ | – |
| *Gpr35* | Class A Orphans | C-HTMR/Heat (*Mrgpra3*$^+$, *Sst*$^+$, *Mrgprd*$^+$), CGRP-γ (*Adra2a*$^+$), C-Heat (*Sstr2*$^+$), CGRP-ε (*Oprk1*$^+$), C-LTMRs | – | Co-expressing CGRP |
| *Gpr4* | Class A Orphans | C-HTMR/Heat (*Sst*$^+$) | $G_s$, $G_{q/11}$, $G_{12/13}$ | – |
| *Gpr45* | Class A Orphans | C-Heat (*Sstr2*$^+$), C-HTMR/Heat (*Sst*$^+$, *Mrgpra3*$^+$, *Mrgprd*$^+$), C-Cold (*Trpm8*$^+$), CGRP-ε (*Oprk1*$^+$), CGRP-γ (*Adra2a*$^+$), Aδ-HTMRs (*Bmpr1b*$^+$) | – | Not expressed |
| *Grm5* | Metabotropic glutamate receptors | CGRP-ε (*Oprk1*$^+$), Aδ-HTMRs (*Bmpr1b*$^+$), CGRP-γ (*Adra2a*$^+$), C-HTMR/Heat (*Mrgpra3*$^+$) | $G_{q/11}$, secondary: $G_s$, $G_{i/o}$ | – |
| *Grm7* | Metabotropic glutamate receptors | C-Heat (*Sstr2*$^+$), C-Cold (*Trpm8*$^+$), CGRP-γ (*Adra2a*$^+$), CGRP-ε (*Oprk1*$^+$), Aδ-HTMRs (*Bmpr1b*$^+$), C-HTMR/Heat (*Mrgprd*$^+$, *Mrgpra3*$^+$) | $G_{i/o}$ | Broadly expressed |
| *Hcrtr1* | Orexin receptors | Aδ-HTMRs (*Bmpr1b*$^+$), C-Heat (*Sstr2*$^+$), CGRP-ε (*Oprk1*$^+$), CGRP-γ (*Adra2a*$^+$) | $G_{q/11}$ | – |
| *Hrh1* | Histamine receptors | C-HTMR/Heat (*Mrgpra3*$^+$, Sst+), CGRP-γ (*Adra2a*$^+$) | $G_{q/11}$ | – |
| *Hrh2* | Histamine receptors | C-HTMR/Heat (*Sst*$^+$, *Mrgpra3*$^+$) | $G_{q/11}$, secondary: $G_s$ | – |
| *Htr1a* | 5-Hydroxytryptamine receptors | C-HTMR/Heat (*Sst*$^+$), CGRP-ε (*Oprk1*$^+$) | $G_{i/o}$ | Not expressed |
| *Htr1b* | 5-Hydroxytryptamine receptors | C-Heat (*Sstr2*$^+$), Aδ-HTMRs (*Bmpr1b*$^+$), Aδ-HTMR/Heat (*Smr2*$^+$), CGRP-γ (*Adra2a*$^+$), CGRP-ε (*Oprk1*$^+$) | $G_{i/o}$ | Co-expressing CGRP |
| *Htr1f* | 5-Hydroxytryptamine receptors | C-HTMR/Heat (*Sst*$^+$) | $G_{i/o}$ | Co-expressing CGRP but not *SST* |
| *Mas1* | Class A Orphans | Aδ-HTMRs (*Bmpr1b*$^+$) | $G_{i/o}$, $G_{q/11}$ | – |

| | | | | |
|---|---|---|---|---|
| *Mrgpra1*[b] | Class A Orphans | C-HTMR/Heat (*Mrgpra3*$^+$), CGRP-ε (*Oprk1*$^+$) | $G_{q/11}$ | – |
| *Mrgpra2a*[b] | Class A Orphans | C-HTMR/Heat (*Mrgpra3*$^+$, *Mrgprd*$^+$, *Sst*$^+$), CGRP-ε (*Oprk1*$^+$), CGRP-γ (*Adra2a*$^+$) | $G_{q/11}$ | – |
| *Mrgpra2b*[b] | Class A Orphans | C-HTMR/Heat (*Mrgpra3*$^+$, *Mrgprd*$^+$, *Sst*$^+$), CGRP-ε (*Oprk1*$^+$), CGRP-γ (*Adra2a*$^+$) | $G_{q/11}$ | – |
| *Mrgpra3*[b] | Class A Orphans | C-HTMR/Heat (*Mrgpra3*$^+$) | $G_{q/11}$ | – |
| *Mrgpra4*[b] | Class A Orphans | C-HTMR/Heat (*Mrgpra3*$^+$) | $G_{q/11}$ | – |
| *Mrgprb5*[b] | Class A Orphans | C-HTMR/Heat (*Mrgpra3*$^+$, *Mrgprd*$^+$) | $G_{q/11}$ | – |
| *Mrgprx1*[b] | Class A Orphans | C-HTMR/Heat (*Mrgpra3*$^+$), CGRP-ε (*Oprk1*$^+$) | $G_{q/11}$ | – |
| *Npffr1* | Neuropeptide FF/neuropeptide AF receptors | CGRP-γ (*Adra2a*$^+$), C-Heat (*Sstr2*$^+$), Aδ-HTMRs (*Bmpr1b*$^+$), C-Cold (*Trpm8*$^+$) | $G_{i/o}$ | – |
| *Npy1r* | Neuropeptide Y receptors | C-Heat (*Sstr2*$^+$), CGRP-γ (*Adra2a*$^+$), CGRP-ε (*Oprk1*$^+$) | $G_{i/o}$ | – |
| *Npy2r* | Neuropeptide Y receptors | C-HTMR/Heat (*Sst*$^+$), Aδ-HTMRs (*Bmpr1b*$^+$), CGRP-γ (*Adra2a*$^+$), CGRP-ε (*Oprk1*$^+$) | $G_{i/o}$, secondary: $G_{q/11}$ | Not expressed |
| *Ntsr2* | Neurotensin receptors | C-Heat (*Sstr2*$^+$), C-Cold (*Trpm8*$^+$), CGRP-ε (*Oprk1*$^+$), CGRP-γ (*Adra2a*$^+$) | $G_{q/11}$ | – |
| *Opn3* | Opsin receptors | C-Heat (*Sstr2*$^+$), CGRP-ε (*Oprk1*$^+$), CGRP-γ (*Adra2a*$^+$) | – | – |
| *Oprk1* | Opioid receptors | CGRP-ε (*Oprk1*$^+$) | $G_{i/o}$, secondary: $G_{12/13}$ | – |
| *Oprm1* | Opioid receptors | C-Heat (*Sstr2*$^+$), C-Cold (*Trpm8*$^+$), C-HTMR/Heat (*Sst*$^+$, *Mrgpra3*$^+$), CGRP-ε (*Oprk1*$^+$), CGRP-γ (*Adra2a*$^+$) | $G_{i/o}$, secondary: $G_{q/11}$ | – |
| *P2ry2* | P2Y receptors | C-HTMR/Heat (*Mrgpra3*$^+$), Aδ-HTMRs (*Bmpr1b*$^+$), CGRP-γ (*Adra2a*$^+$), C-Heat (*Sstr2*$^+$), Aδ-HTMR/Heat (*Smr2*$^+$) | $G_{q/11}$, secondary: $G_{i/o}$, $G_{12/13}$ | – |
| *Prokr2* | Prokineticin receptors | Aδ-HTMRs (*Bmpr1b*$^+$), CGRP-γ (*Adra2a*$^+$) | $G_{q/11}$, secondary: $G_s$, $G_{i/o}$ | – |
| *Ptafr* | Platelet-activating factor receptor | C-HTMR/Heat (*Sst*$^+$) | $G_{i/o}$, $G_{q/11}$ | – |
| *Ptgdr* | Prostanoid receptors | C-Heat (*Sstr2*$^+$), C-HTMR/Heat (*Mrgprd*$^+$, *Mrgpra3*$^+$) | $G_s$ | – |
| *Ptger1* | Prostanoid receptors | CGRP-ε (*Oprk1*$^+$), C-Heat (*Sstr2*$^+$), CGRP-γ (*Adra2a*$^+$), C-HTMR/Heat (*Sst*$^+$) | $G_{q/11}$, secondary: $G_{i/o}$ | – |
| *Ptgir* | Prostanoid receptors | CGRP-ε (*Oprk1*$^+$), CGRP-γ (*Adra2a*$^+$), Aδ-HTMRs (*Bmpr1b*$^+$), C-Heat (*Sstr2*$^+$), Aδ-HTMR/Heat (*Smr2*$^+$), C-HTMR/Heat (*Mrgpra3*$^+$, *Sst*$^+$) | $G_s$, secondary: $G_{i/o}$, $G_{q/11}$ | – |

| | | | | |
|---|---|---|---|---|
| *Rho* | Opsin receptors | C-Heat (*Sstr2*$^+$), CGRP-ε (*Oprk1*$^+$), CGRP-γ (*Adra2a*$^+$), Aδ-HTMRs (*Bmpr1b*$^+$) | – | – |
| *Rxfp3* | Relaxin family peptide receptors | CGRP-γ (*Adra2a*$^+$) | $G_{i/o}$ | – |
| *S1pr1* | Lysophospholipid (S1P) receptors | C-HTMR/Heat (*Sst*$^+$), CGRP-ε (*Oprk1*$^+$), C-Heat (*Sstr2*$^+$), CGRP-γ (*Adra2a*$^+$) | $G_{i/o}$ | – |
| *S1pr2* | Lysophospholipid (S1P) receptors | C-HTMR/Heat (*Sst*$^+$, *Mrgpra3*$^+$), C-Heat (*Sstr2*$^+$) | $G_s$, $G_{q/11}$, $G_{12/13}$ | – |
| *S1pr3* | Lysophospholipid (S1P) receptors | C-Heat (*Sstr2*$^+$), CGRP-ε (*Oprk1*$^+$), CGRP-γ (*Adra2a*$^+$), Aδ-HTMRs (*Bmpr1b*$^+$) | $G_{i/o}$, $G_{q/11}$, $G_{12/13}$ | – |
| *Sstr1* | Somatostatin receptors | CGRP-γ (*Adra2a*$^+$), CGRP-ε (*Oprk1*$^+$) | $G_{i/o}$ | – |
| *Sstr2* | Somatostatin receptors | C-Heat (*Sstr2*$^+$) | $G_{i/o}$ | Co-expressing CGRP |
| *Tbxa2r* | Prostanoid receptors | CGRP-γ (*Adra2a*$^+$) | $G_{q/11}$ | – |
| *Vmn1r85*[c] | Vomeronasal receptors | CGRP-ε (*Oprk1*$^+$), CGRP-γ (*Adra2a*$^+$) | – | – |
| *Vmn1r89*[c] | Vomeronasal receptors | CGRP-ε (*Oprk1*$^+$) | – | – |
| *Vmn2r113*[c] | Vomeronasal receptors | C-HTMR/Heat (*Mrgpra3*$^+$) | – | – |

**Table S1. List of GPCRs that are selectively expressed in CGRP$^+$ sensory neurons, related to Figure 1.** The GPCRs that are restrictively expressed in one or more CGRP$^+$ DRG sensory neuron subtypes were identified by mapping individual genes to our single cell RNA-seq dataset [5]. Expression patterns are listed. Information on the transduction pathways were obtained from the IUPHAR/BPS Guide to PHARMACOLOGY. Expression patterns in human DRGs inferred from the RNAscope data are also included. [a] Human ortholog *GPR165P* is a pseudogene. [b] Not well conserved between mouse and human. [c] No human ortholog.
